# Polyamine biosynthesis and eIF5A hypusination are modulated by the DNA tumor virus KSHV and promote KSHV viral infection

**DOI:** 10.1101/2020.12.19.423609

**Authors:** Guillaume N. Fiches, Ayan Biswas, Dawei Zhou, Weili Kong, Maxime Jean, Netty G. Santoso, Jian Zhu

**Affiliations:** Department of Pathology, Ohio State University College of Medicine, Columbus, Ohio, 43210, United States; Gladstone Institute of Virology and Immunology, University of California, San Francisco, California, 94158, United States; Department of Neurology, University of Rochester Medical center, Rochester NY 14642, USA

**Author notes:** To whom correspondence should be addressed: Jian Zhu.

**Keywords:** Polyamine, KSHV, tumor virus, hypusine, eIF5A, lytic replication, latency, ORF50/RTA, ORF73/LANA, ODC1, spermidine

## Abstract

Polyamines are critical metabolites involved in various cellular processes and often dysregulated in cancers. Kaposi’s sarcoma associated Herpesvirus (KSHV) is a defined oncogenic virus belonging to the sub-family of human gamma-herpesviruses. KSHV infection leads to the profound alteration of host metabolic landscape to favor the development of KSHV-associated malignancies. In our studies, we identified that polyamine biosynthesis and eIF5A hypusination are dynamically regulated by KSHV infection likely through the modulation of key enzymes of these pathways, such as ODC1, and that in return these metabolic pathways are required for both KSHV lytic switch from latency and *de novo* infection. The further analysis unraveled that translation of critical KSHV latent and lytic proteins (LANA, RTA) depends on eIF5A hypusination. We also demonstrated that KSHV infection can be efficiently and specifically suppressed by using inhibitors targeting either polyamine biosynthesis or eIF5A hypusination. Above all, our results illustrated that the dynamic and profound interaction of a DNA tumor virus (KSHV) with host polyamine biosynthesis and eIF5A hypusination metabolic pathways promote viral propagation and oncogenesis, which serve as new therapeutic targets to treat KSHV-associated malignancies.

## Introduction

Polyamines, namely putrescine, spermidine and spermine, are low-molecular weight metabolites that are ubiquitous in eukaryotic life. Present in cells at millimolar concentrations, these polycations effectively binds DNA, RNA, and phospholipids thanks to their positive charge at physiological conditions [1] and, in turn, regulate many cellular processes, such as transcription, translation, cell cycle, chromatin remodeling and autophagy [2–7]. Consequently, their metabolism is tightly regulated by multiple layers of feedback loops and interconversion mechanisms [8, 9]. Ornithine decarboxylase 1 (ODC1) is the rate-limiting enzyme that modulates the initial step of the polyamine biosynthesis pathway by converting ornithine into putrescine. Putrescine is then converted into spermidine by the spermidine synthase (SRM) which is subsequently converted into spermine by the spermine synthase (SMS). In addition, catabolic enzymes, such as spermidine-spermine N1-acetyltransferase (SSAT-1), polyamine oxidase (PAO), and spermine oxidase (SMOX), can acetylate or oxidize polyamines to mediate their export and/or convert back to previous stages.

However, the homeostatic balance of polyamines is often disrupted in cancers. Indeed, upregulated metabolism of polyamines has been observed in cancers in order to fulfill the increased needs associated with tumorigenesis and rapid growth of tumor cells (reviewed in [10, 11]). For instance, polyamines, particularly spermidine, are required for the hypusination of the eukaryotic initiation factor 5A (eIF5A), which has been found upregulated in multiple types of cancers [12–14]. Hypusination is a unique post-translational modification, only reported to occur on eIF5A so far [15, 16], which enables the selective control of protein translation through a conserved biochemical mechanism targeting “hard-to-translate” region, such as polyproline stretches [17, 18]. Consequently, eIF5A hypusination has emerged as a key regulator of the polyamines’ downstream cellular processes, such as autophagy [19, 20].

Furthermore, investigation of polyamines’ role in regulating viral infections started nearly 50 years ago. Earlier reports suggested that polyamines benefit the viral packaging of large DNA viruses, such as herpes simplex virus (HSV-1) [21], Vaccinia virus (VACV) [22], and polioviruses [23]. Only very recently, it has been reported that polyamines are involved in the viral life cycle of RNA viruses to promote transcription, translation, and viral packaging, including Zika and Chikungunya viruses [24], Ebola virus [25, 26], Rift valley fever and LaCrosse viruses [27, 28]. In return, viruses often manipulate the polyamine pathway, illustrated by herpesviruses. Human cytomegalovirus (HCMV) is known to induce the expression of ODC1 [29], while ODC1 inhibition using its inhibitor DFMO limits viral replication of HCMV [30]. HSV-1 upregulates the expression of S-adenosyl methionine decarboxylase (SAMDC) [31], while Epstein-Barr virus (EBV) downregulates SSAT-1 [32].

In contrast, there is almost no knowledge regarding to the functional contribution of polyamines to Kaposi’s sarcoma associated Herpesvirus (KSHV), a relatively recently discovered herpesvirus. KSHV is a human γ-herpesvirus and the etiological agent leading to Kaposi’s sarcoma (KS) [33, 34] and associated with two lymphoproliferative disorders, primary effusion lymphoma (PEL) [35] and Multicentric Castleman Disease (MCD) [36]. KSHV has the capacity to establish a life-long infection in the infected individual and persist primarily in infected B lymphocytes and endothelial cells in a quiescent state. During viral latency, only a limited subset of viral latent genes is expressed. Latent infection of KSHV supports the maintenance of viral episomes, and also modulates host cellular environment, including regulation of metabolic pathways, to promote cell proliferation [37–39]. Latent KSHV can be reactivated, and expression of viral lytic genes is turned on and new virions are produced during viral lytic cycle. Although KSHV remains latent in most infected cells, which plays a major role in viral tumorigenesis, lytic replication of KSHV is critical for viral dissemination and also contributes to tumor progression [40, 41]. Hence, it is critical to understand how latent KSHV shapes the host cellular environment to accommodate and benefit its lytic reactivation. In the present study, we reported that polyamine metabolism and eIF5A hypusination are modulated by KSHV, which in return promotes KSHV viral infection.

## Results

### KSHV dynamically modulates intracellular polyamines

We first analyzed the global polyamine level in KSHV latently infected tumor cells and also upon lytic reactivation. The renal carcinoma cell line with epithelial-cell origin, known as SLK, and its counterpart cell line latently infected with KSHV BAC16 strain as well as transduced with doxycycline (Dox) inducible RTA, named iSLK.BAC16 [42], were treated with either Dox to induce KSHV RTA expression and consequently lytic reactivation, or vehicle control to keep KSHV at latency. Intracellular total polyamines were stained by using the PolyamineRED reagent [43] and visualized via confocal microscopy. We observed that the total polyamine level was significantly higher in iSLK.BAC16 vs SLK cells without Dox (latent state) (**Fig 1A,B**). However, intracellular polyamines markedly and specifically decreased during lytic replication in Dox-treated iSLK.BAC16 but remained unchanged in the dox-treated naive SLK cells (**Fig 1A,B**). We further analyzed the free, major polyamine species in these cells via thin-layer chromatography (TLC), including putrescine (put), spermidine (spd), and spermine (spm) (**Fig 1C,D**). Putrescine remained relatively stable in both SLK and iSLK.BAC16 cells treated with vehicle control or Dox (**Fig 1C,D, S1A**). Spermine was slightly increased in iSLK.BAC16 vs SLK cells without Dox, while Dox moderately reduced spermine in iSLK.BAC16 cells (**Fig 1C,D, S1C**). On the contrary, spermidine was significantly decreased in iSLK.BAC16 cells at both -/+ Dox conditions (38.3% and 68.6% reduction respectively) (**Fig 1C,D, S1B**). In addition, we also calculated the relative fraction of polyamine species, which revealed that the proportion of spermidine was the most impacted by KSHV latency and lytic reactivation. Spermidine fraction in iSLK.BAC16 cells was indeed lesser than SLK cells at both -/+ Dox conditions (without Dox: 27.6% *vs* 39.9%; with Dox: 19.1% *vs* 39.6%), and Dox particularly led to the further decrease of spermidine fraction in iSLK.BAC16 cells but not SLK cells (**Fig 1E**). KSHV latency led to a slight increase of spermine fraction (**Fig 1D,E**). However, spermine fraction decreased due to KSHV lytic induction, while putrescine fraction remained stable (**Fig 1D**) resulting in an increase of the relative portion of putrescine (**Fig 1E**). We also performed the above PolyamineRED staining and TLC assays for a KSHV-positive lymphoma cell line transduced with Dox-inducible RTA, TREx BCBL1-RTA, and a KSHV-negative counterpart, BJAB. Similarly, we noticed that the total polyamine level was significantly higher in TREx BCBL1-RTA vs BJAB cells without Dox and that it markedly decreased during lytic replication in Dox-treated TREx BCBL1-RTA cells but not in Dox-treated naïve BJAB (**Fig S1D,E**). TLC assay also confirmed that spermidine was the most reduced polyamine species in Dox-treated TREx BCBL1-RTA cells (**Figure S1F,G**).

**Figure 1.**
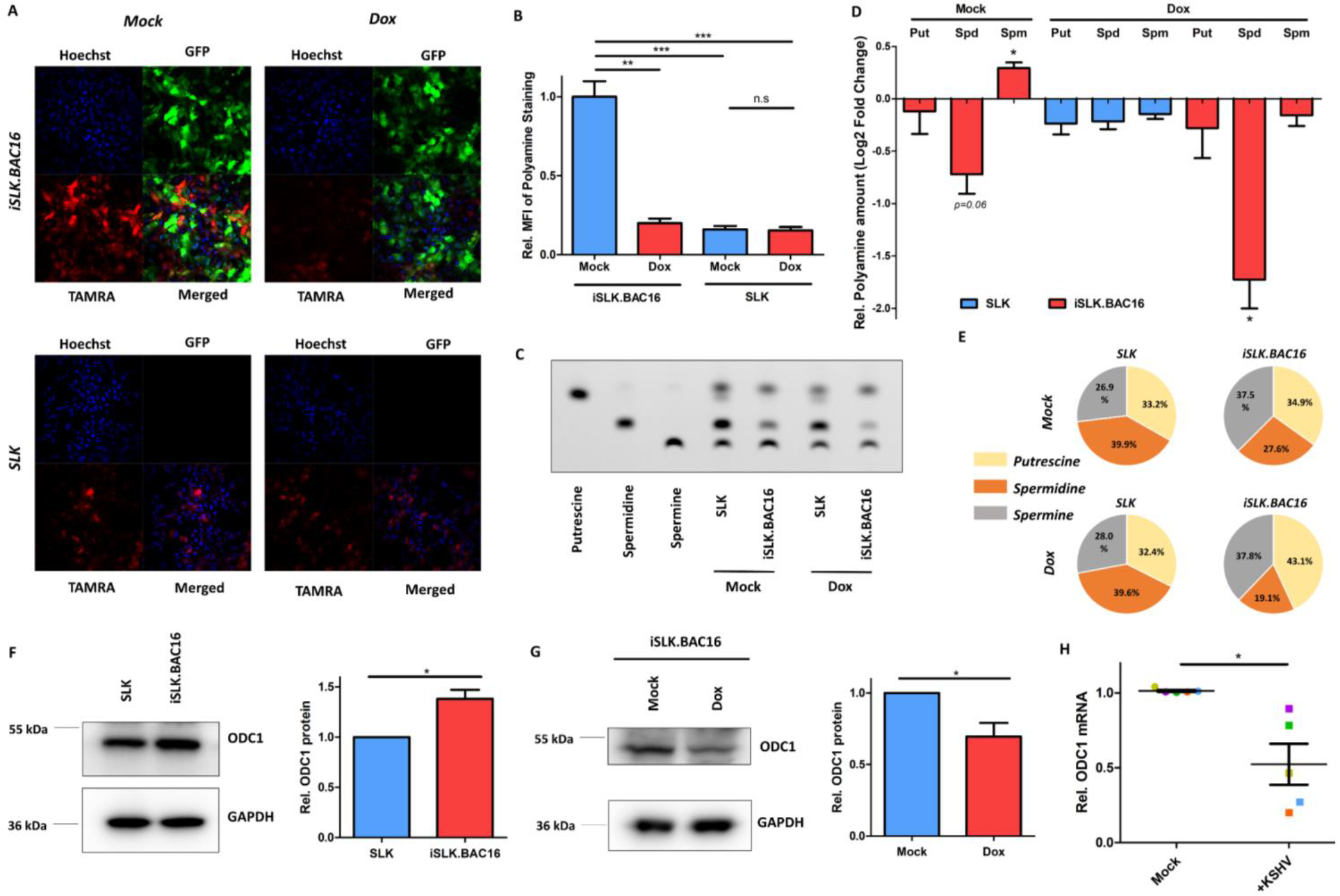
KSHV dynamically modulates the level of intracellular polyamines. (A, B). Intracellular polyamines in SLK and iSLK.BAC16 cells treated with doxycycline (Dox; 1μg/mL) for 48h or mock were stained with PolyamineRED (TAMRA) and visualized by confocal fluorescence microscopy (A), whereas nuclei were labelled with Hoechst (Blue). GFP is constitutively expressed in cells infected with KSHV BAC16. Mean fluorescence intensity (MFI) of polyamine fluorescence signal (TAMRA) of ≥ 1500 cells (B) was determined and normalized to mock-treated iSLK.BAC16 cells. (C-E). Intracellular polyamine species (putrescine [Put], spermidine [Spd], spermine [Spm]) in SLK and iSLK.BAC16 cells treated with Dox for 48h or mock were analyzed by thin-layer chromatography (TLC) with pure individual polyamine species used as reference (C). Relative changes of polyamine species (Put, Spd, Spm) in TLC results were determined and normalized to mock-treated SLK cells (D), or displayed as the relative proportion of total intracellular polyamines in pie chart (E). (F, G). ODC1 protein in SLK and iSLK.BAC16 cells (F) as well as in iSLK.BAC16 cells treated with Dox or mock (G) was measured by immunoblotting. Intensity of protein bands was determined by using AlphaView SA (software) and normalized to GAPDH. (H) Primary tonsillar B lymphocytes from 5 healthy donors were isolated and infected with KSHV BAC16 viruses or mock treated. mRNA level of ODC1 in these cells at 3 dpi was measured by RT-qPCR assays and normalized to mock treatment. Results were calculated from n=3-4 independent experiments and presented as mean ± SEM (* p<0.05; ** p<0.01; *** p<0.001, two-tailed paired Student t-test).

Taken together, these results clearly showed that KSHV infection distinctly and dynamically modulates the level of intracellular polyamines at latent and lytic phases. It has been reported that c-Myc is upregulated by LANA during KSHV latency [44], while ODC1 is transcriptionally activated by c-Myc [45, 46]. Indeed, we found that ODC1 is slightly upregulated in iSLK.BAC16 vs SLK cells (**Fig 1F**). In contrast, KSHV lytic induction resulted in the decrease of ODC1 protein in Dox-treated iSLK.BAC16 (**Fig 1G**), likely because c-Myc is silenced due to KSHV lytic reactivation [47, 48]. Consistently, we also observed that ODC1 mRNA level also decreased at 3 dpi in primary tonsillar B cells isolated from healthy donors, known to be susceptible to KSHV infection and maintain active lytic replication [49], which were spinoculated with KSHV.BAC16 viruses, followed by RT-qPCR analysis (**Fig 1H**).

### Polyamine synthesis enzymes are required for KSHV lytic reactivation

We sought to further determine the contribution of the polyamine pathway to KSHV infection by investigating several key synthesis enzymes from this pathway (**Fig 2A**). ODC1 is critical to catalyze the initial step of polyamine synthesis by converting ornithine into putrescine. HEK293 cells harboring a recombinant KSHV r219 strain (HEK293.r219) were transiently transfected with siRNAs targeting ODC1 or non-targeting (NT) control, and ODC1 knockdown was verified by qPCR and immunoblotting assays (**Fig 2B,C**). TLC assays showed that ODC1 knockdown led to the moderate reduction of putrescine and spermine, by 23% and 16% respectively, and the drastic depletion of spermidine by 85% reduction, averaged from three ODC1 siRNAs **(Figure 2D,E**). KSHV r219 strain carries GFP and RFP fluorescent reporter genes allowing visualization of KSHV latent and lytic infection respectively [50]. KSHV lytic reactivation was induced by treating HEK293.r219 cells with 12-O-Tetradecanoylphorbol 13-acetate and Sodium Butyrate (TPA+NaB). Fluorescence imaging indicated that TPA+NaB-induced RFP expression was significantly reduced due to ODC1 knockdown for all three ODC1 siRNAs (**Fig 2F**). qPCR assays consistently showed that ODC1 knockdown by its three siRNAs significantly reduces the TPA+NaB-induced expression of KSHV lytic genes, including ORF50/RTA, K8/K-bZIP, and ORF26 genes, respectively representing immediate early, early and late lytic genes (**Fig 2G**), which is supported by the immunoblotting assays confirming the reduced protein level of KSHV lytic genes, ORF45 and K8.1A/B, in HEK293.r219 cells transfected with ODC1 siRNAs (**Fig 2H**). In addition, ODC1 knockdown even reduced the residual expression of KSHV lytic genes that occurs during spontaneous lytic replication (**Fig S2A**). The similar effect of ODC1 knockdown on KSHV lytic reactivation was observed in HEK293.r219 cells, in which KSHV lytic reactivation was alternatively induced by ectopic expression of KSHV ORF50 cDNA (**Fig S2B,C**), as well as in iSLK.BAC16 cells treated with Dox+NaB to induce reactivation of latent KSHV (**Fig S2D,E**). Taken together, data demonstrated the presence of ODC1 is required for the efficient KSHV lytic reactivation.

**Figure 2.**
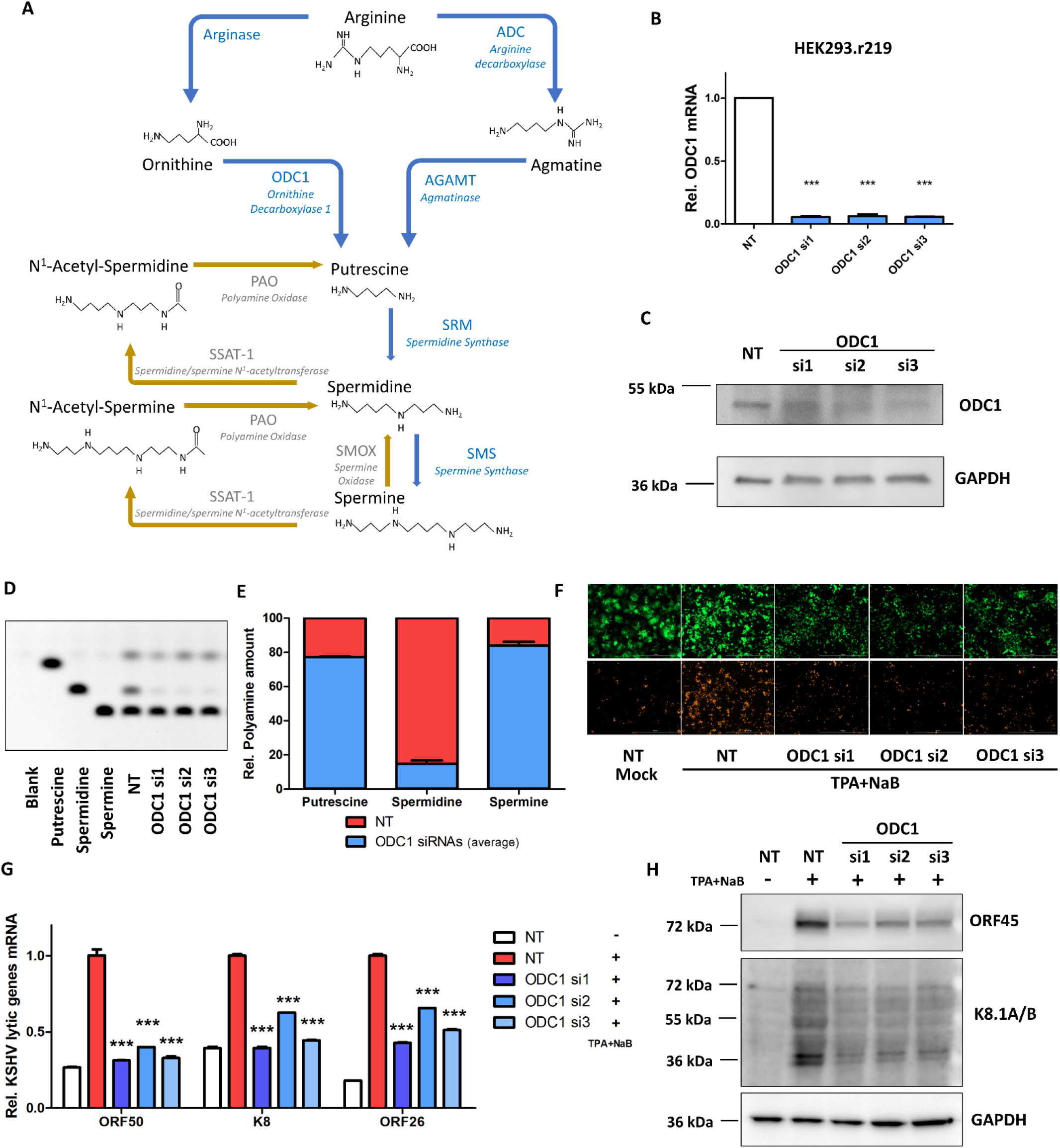
ODC1 is required for KSHV lytic reactivation. (A). Schematic view of the polyamine biosynthesis pathway. (B, C). ODC1 knockdown in HEK293.r219 cells transfected with ODC1 siRNAs (si1, si2, si3) or non-targeting control siRNA (NT) was analyzed by RT-qPCR (B) or immunoblotting (C). (D, E). Intracellular polyamine species (Put, Spd, Spm) in HEK293.r219 cells transfected with ODC1 siRNAs (si1, si2, si3) or NT were analyzed by TLC with pure individual polyamine species used as reference (D). Relative polyamine species (Put, Spd, Spm) in above cells were quantified (average of ODC1 si1-3) and normalized to NT-transfected cells. Results were calculated from n=2 independent repeats. (F-H). HEK293.r219 cells transfected with ODC1 siRNAs (si1, si2, si3) or NT were treated with TPA (20 ng/mL) + NaB (0.3mM) for 48h, and visualized by fluorescence imaging (F). Expression of RFP protein indicates KSHV lytic reactivation, while GFP signal means that cells are KSHV-infected. mRNA level of KSHV lytic genes (ORF50/RTA, K8/K-bZIP, ORF26) in above cells was analyzed by RT-qPCR assays and normalized to NT-transfected induced cells (G). Protein level of KSHV lytic genes (ORF45, K8.1A/B) was analyzed by immunoblotting assays (H). GAPDH was used as the loading control. Results were calculated from n=3 independent experiments and presented as mean ± SEM (* p<0.05; ** p<0.01; *** p<0.001, two-tailed paired Student t-test).

Using the similar approaches, we also determined the relevance of other polyamine synthesis enzymes to KSHV lytic reactivation. Beside ODC1, agmatinase (AGMAT) can convert agmatine to putrescine as well (**Fig 2A**). Although its importance in the polyamine metabolism was initially described for the lower organisms [51], AGMAT has only been recently studied for mammals [52]. Expression of AGMAT was significantly reduced by its two siRNAs in HEK293.r219 (**Fig 3A**), which caused the reduction of TPA+NaB-induced expression of RFP (**Fig 3B**) and KSHV lytic genes (**Fig 3C**) measured by fluorescence imaging and qPCR assays respectively. Putrescine is the shortest biogenic polyamine and converted to spermidine by the spermidine synthase (SRM), which is subsequently converted to spermine by the spermine synthase (SMS) (**Fig 2A**). Two siRNAs targeting SRM or SMS successfully led to their knockdown (**Fig 3D,E**), which also caused the reduction of TPA+NaB-induced expression of RFP **(Fig 3F)** and KSHV lytic genes **(Fig 3G,H)**measured by fluorescence imaging and qPCR assays respectively. SRM or SMS knockdown also reduced the residual expression of KSHV lytic genes that occurs during spontaneous lytic replication (**Fig S3A,B**). In addition, spermidine-spermine N1-acetyltransferase (SSAT-1) is a key catabolic enzyme in the polyamine pathway that acetylates spermidine and spermine, resulting in either their export out of the cell or their conversion by polyamine oxidase (PAO) to putrescine and spermidine respectively. Ribavirin is a FDA-approved drug that has been recently shown to induce SSAT-1 and thus deplete intracellular polyamines [53]. Our results identified that ribavirin severely impairs the Dox-induced KSHV lytic gene expression (**Fig 3I)** and viral genome amplification **(Fig 3J**) in iSLK.BAC16 cells in a dose-dependent manner which was sufficient to increase SSAT-1 protein level (**Fig S3C**) with limited cytotoxicity (**Fig S3D**) in these cells. Collectively, our investigation of several key polyamine synthesis enzymes undoubtedly verified that polyamine homeostasis is critical to KSHV lytic reactivation.

**Figure 3.**
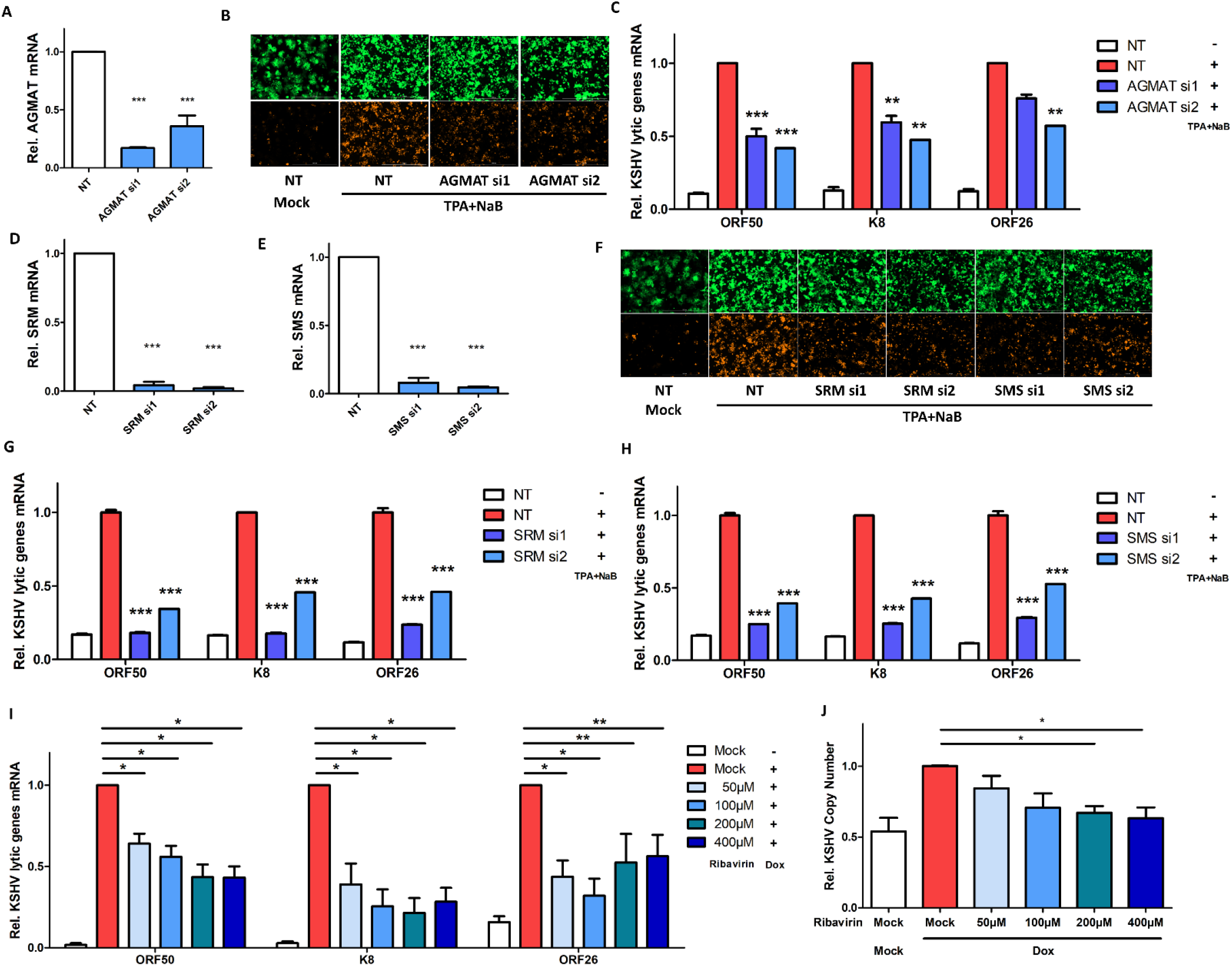
Other polyamine enzymes contribute to KSHV lytic reactivation. (A-C) AGMAT knockdown in HEK293.r219 cells transfected with AGMAT siRNAs (si1, si2) or NT was analyzed by RT-qPCR assays (A). HEK293.r219 cells transfected with AGMAT siRNAs (si1, si2) or NT was treated with TPA (20 ng/mL) + NaB (0.3mM) for 48h, and visualized by fluorescence imaging (B). mRNA level of KSHV lytic genes (ORF50/RTA, K8/K-bZIP, ORF26) in above cells was analyzed by RT-qPCR assays and normalized to NT-transfected induced cells (C). (D, E). SRM (D) or SMS (E) knockdown in HEK293.r219 cells transfected with their siRNAs (si1, si2) or NT was analyzed by RT-qPCR assays. (F-H). HEK293.r219 cells transfected with SRM/SMS siRNAs (si1, si2) or NT was treated with TPA (20 ng/mL) + NaB (0.3mM) for 48h, and visualized by fluorescence imaging (F). mRNA level of KSHV lytic genes (ORF50/RTA, K8/K-bZIP, ORF26) in SRM (G) or SMS (H) depleted, TPA+NaB-induced HEK293.r219 cells was analyzed by RT-qPCR assays and normalized to NT-transfected induced cells. (I, J). mRNA level of KSHV lytic genes (ORF50/RTA, K8/K-bZIP, ORF26) in iSLK.BAC16 cells pretreated with increasing doses of ribavirin for 24h and subsequently induced with Dox (1μg/mL) was analyzed by RT-qPCR assays (I). Copy number of KSHV genomes in above cells was also analyzed (J). Results were calculated from n=3 independent experiments and presented as mean ± SEM (* p<0.05; ** p<0.01; *** p<0.001, two-tailed paired Student t-test).

### Inhibition of polyamine synthesis efficiently blocks KSHV lytic reactivation

Inhibitors have been developed to target specific enzymes or steps of polyamine pathway for treating various diseases, especially cancers. α-difluoromethylornithine (DFMO) is a well-tolerated, potent, and irreversible inhibitor of ODC1. Despite its poor pharmacokinetics, DFMO has significant therapeutic potential as a FDA-approved drug for treating trypanosoma (African sleeping sickness) [54], and it also has shown promising results in the on-going clinical trial for treating high-risk neuroblastoma, a severe form of pediatric tumor [ClinicalTrials.gov Identifier: NCT02679144] [55–57]. In our studies, DFMO was used as a chemical probe to inhibit ODC1 function and block downstream polyamine synthesis. Following 24hr treatment with DFMO, HEK293.r219 cells were transfected with an ectopic KSHV ORF50 cDNA to reactivate latent KSHV. DFMO reduced the RFP signal in a dose-dependent manner (**Fig 4A**). The qPCR assays of these cells also showed that DFMO significantly decreases expression of KSHV K8/bZIP early lytic gene and the copy number of KSHV genomes with comparable IC_50_ (drug concentration for 50% of maximal inhibitory effect) of 95.9μM and 103.2μM respectively (**Fig 4B**). DFMO also decreased the expression of KSHV latent genes (ORF71/v-FLIP, ORF72/v-Cyclin, and ORF73/LANA with IC_50_ = 103.2μM, 115.6μM, 210.8μM respectively) in these cells as well. To confirm that DFMO’s anti-KSHV effect is through depletion of polyamines, we also performed the rescue experiments by adding exogenous polyamines in the form of polyamines mixture (PA Supp), spermidine (Spd), or spermine (Spm) to the DFMO-treated (500μM ≈ IC_90_), ORF50-transfected HEK293.r219 cells (**Fig 4C**), which was able to restore the DMFO-inhibited lytic reactivation of KSHV through measurement of RFP-positive cells as well as expression of KSHV lytic genes (K8, ORF26) but caused no obvious cytotoxicity (**Fig S4A**). Likewise, DFMO inhibited the Dox+NaB-induced KSHV reactivation in iSLK.BAC16 cells as shown through qPCR analysis of copy number of KSHV genomes and KSHV lytic gene expression and immunoblottings (**Fig 4D,E**). Similar effect of DFMO was also confirmed in Dox-treated TREx BCBL1-RTA cells by immunoblottings of KSHV lytic proteins (**Fig S4B**).

**Figure 4.**
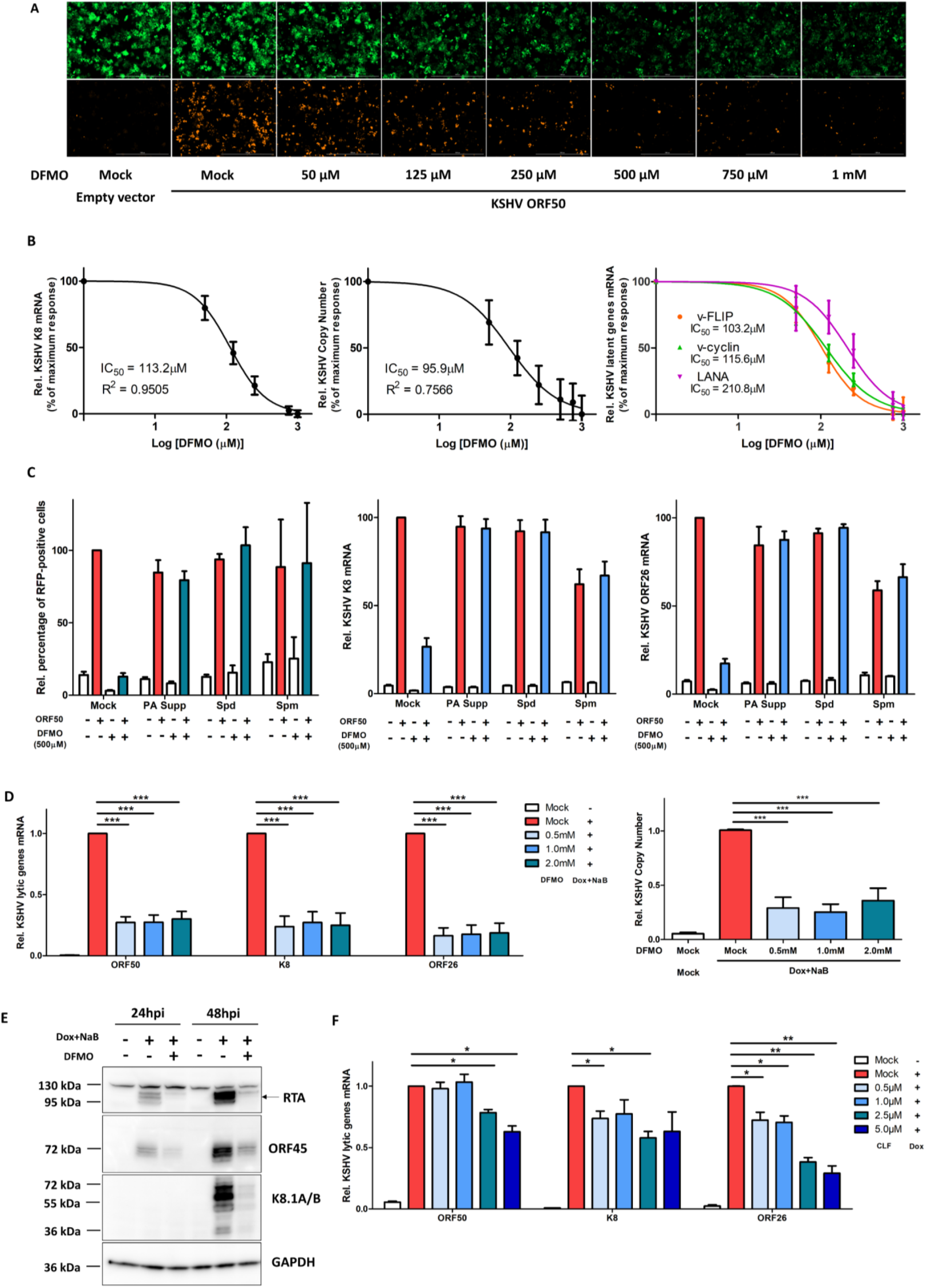
Polyamine depletion efficiently blocks KSHV lytic reactivation. (A, B). HEK293.r219 cells pretreated with progressively increasing doses of 2-difluoromethylornithine (DFMO) for 24h were subsequently induced by ectopic expression of ORF50 for 48h or left un-induced using the empty vector. Above cells were then visualized by fluorescence imaging (A). mRNA level of KSHV K8/K-bZIP (left panel) and latent genes (ORF71/v-FLIP, ORF72/v-Cyclin, and ORF73/LANA; right panel) in above cells were analyzed by RT-qPCR assays and normalized to mock-treated induced cells. The drug effect at a series of doses was plotted as percentage of maximum response by using GraphPad PRISM 5. Relative copy number of KSHV genomes (middle panel) was also analyzed. IC_50_ was determined for DFMO’s inhibitory effect. (C). Exogenous polyamines (mixed polyamines supplement [PA Supp, 5x], Spermidine [Spd, 10μM] or Spermine [Spm, 10μM]) were complemented into the culture media of HEK293.r219 cells treated with DFMO (500μM) and induced by ectopic expression of ORF50 for 48h or left un-induced using the empty vector. The above cells were labelled with Hoechst and visualized by fluorescence imaging to determine percentage of RFP-positive cells (left panel). mRNA level of KSHV lytic genes (K8/K-bZIP [middle panel], ORF26 [right panel]) in above cells were analyzed by RT-qPCR assays. All results were normalized to mock-treated ORF50-induced cells. (D, E). mRNA level of KSHV lytic genes (ORF50/RTA, K8/K-bZIP and ORF26) in iSLK.BAC16 cells pre-treated with increasing doses of DFMO for 24h and subsequently induced by Dox (1μg/mL) + NaB (1mM) for 48h were analyzed by RT-qPCR assays and normalized to mock-treated induced cells (D, left panel). Relative copy number of KSHV genomes in above cells was also analyzed (D, right panel). Protein level of KSHV lytic genes (ORF45, K8.1A/B) was analyzed by immunoblotting (E). GAPDH was used as the loading control. (F). mRNA level of KSHV lytic genes (ORF50/RTA, K8/K-bZIP and ORF26) in iSLK.BAC16 cells pre-treated with increasing doses of clofazimine (CLF) for 24h and subsequently induced by Dox (1μg/mL) for 48h were analyzed by RT-qPCR assays and normalized to mock-treated induced cells. Results were calculated from n=3 independent experiments and presented as mean ± SEM (* p<0.05; ** p<0.01; *** p<0.001, two-tailed paired Student t-test).

Alternatively, we also tested clofazimine (CLF), another FDA-approved drug that has been shown to inhibit ODC1 transcriptional activation [58]. CLF is used for treating drug-resistant tuberculosis [59] and has also demonstrated the potential for treating cancers, such as multiple myeloma [58]. Indeed, CLF treatment led to the reduction of Dox-induced KSHV lytic gene expression in iSLK.BAC16 cells in a dose-dependent manner (**Fig 4F**) without obvious cytotoxicity (**Fig S4C**). Overall, these results clearly demonstrated that polyamine synthesis is required for efficient KSHV lytic reactivation, and that FDA-approved drugs inhibiting the key enzyme ODC1 can be used to block KSHV lytic reactivation and the following viral dissemination.

### eIF5A hypusination is regulated by KSHV and required for its lytic reactivation

Autophagy is a critical downstream cellular process regulated by polyamines, particularly spermidine. Spermidine is a unique substrate of the deoxyhypusine synthase (DHPS) that catalyzes the formation of deoxyhypusinated eIF5A, followed by the deoxyhypusine hydrolase (DOHH) mediated hypusination of eIF5A at K50 [60] (**Fig 5A**). Such post-translational modification is very unique and known to only occur to eIF5A [15, 16]), which has been recently illustrated as critical for autophagy activation [19, 20]. It is also known that KSHV RTA induces autophagy [61] that plays a role in KSHV lytic reactivation, likely through transcriptional regulation of autophagy genes, which has been reported in the case of EBV [62]. Thus, we postulated that spermidine-regulated eIF5A hypusination connects the upstream polyamine pathway with the downstream autophagy activation and would be important to KSHV lytic reactivation, which has never been investigated. To confirm this thought, we transfected the siRNAs targeting DHPS or NT control in iSLK.BAC16 cells. DHPS was efficiently knocked down by its two siRNAs (**Fig 5B**), which led to the significant decrease of Dox-induced KSHV lytic gene expression in these cells (**Fig 5C**).

**Figure 5.**
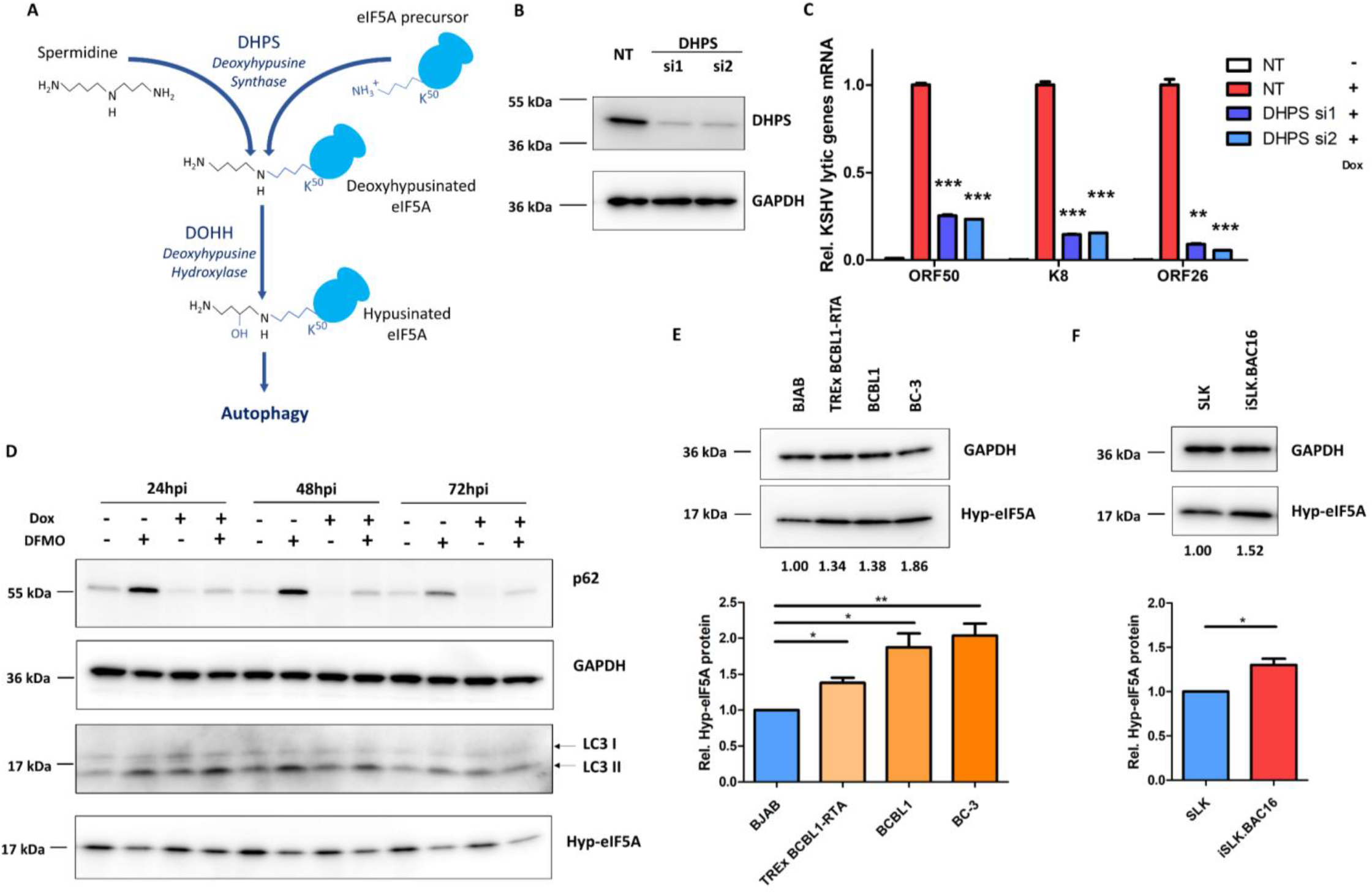
KSHV lytic reactivation depends on eIF5A hypusination. (A). Schematic view of eIF5A hypusination that links spermidine with autophagy activation. (B, C). DHPS knockdown of iSLK.BAC16 cells transfected with DHPS siRNAs (si1, si2) or NT was analyzed by immunoblotting (B). GAPDH was used as a loading control. mRNA level of KSHV lytic genes (ORF50/RTA, K8/K-bZIP and ORF26) in above cells induced with Dox (1μg/mL) was analyzed by RT-qPCR assays and normalized to NT-transfected induced cells (C). (D). Protein level of autophagy markers (p62/SQSTM1, LC3 I/II) as well as hypusine-eIF5A (hyp-eIF5A) in iSLK.BAC16 cells pre-treated with DFMO (500μM) for 24 h and subsequently induced with Dox (1μg/mL) up to 72h was analyzed by immunoblotting assays. GAPDH was used as the loading control. (E, F). Protein level of hyp-eIF5A protein levels in KSHV-infected PEL cells lines (BCBL1 [wild-type], TREx BCBL1-RTA, BC-3) and the KSHV-negative B cell line BJAB (E), or iSLK.BAC16 and SLK cells (F), was analyzed by immunoblotting assays. Intensity of protein bands was determined by using AlphaView SA (software) and normalized to GAPDH. Results were calculated from n=3-4 independent experiments and presented as mean ± SEM (* p<0.05; ** p<0.01; *** p<0.001, two-tailed paired Student t-test).

Reminiscent of earlier studies showing that DFMO blocks autophagy through affecting spermidine [63, 64], we further confirmed that DFMO indeed concurrently reduces the level of hypusinated eIF5A (hyp-eIF5A) as well as impairs autophagy activation assessed by the protein level of p62/SQSTM1 and LC3-I/II while inhibiting KSHV lytic reactivation (**Fig 5D**). Degradation of p62 indicates the lysosomal degradation of autophagosomes and the completion of autophagy events. Protein level of p62 was rapidly depleted in Dox-treated iSLK.BAC16 cells, while DFMO restored p62 in both Dox-treated and un-treated cells. DFMO also led to the accumulation of LC3-II but had no severe impact on LC3I in these cells. We also determined that DFMO blocks upregulation of LC3A and IRGM (immunity-related GTPase family M), another important autophagy related gene [65] often found dysregulated during viral infection [66], due to Dox-induced KSHV reactivation in iSLK.BAC16 cells (**Fig S5A**).

As we noticed that KSHV infection modulates intracellular polyamine level, we were curious whether it is the same case for eIF5A hypusination. We noticed that the protein level of hypusinated eIF5A is consistently higher in all three tested KSHV-positive lymphoma cell lines (TREx BCBL1-RTA, BCBL1, BC-3) comparing to KSHV-negative BJAB cells (**Fig 5E**). It is also true for iSLK.BAC16 in comparison to SLK cells (**Fig 5F**). These results suggested that, beyond the elevated polyamines, the hypusinated eIF5A was also increased due to KSHV latent infection. In addition, treatment of TPA+NaB to induce KSHV lytic reactivation also led to the slight increase of hypusinated eIF5A in HEK293.r219 cells (**Fig S5B**), suggesting that KSHV lytic reactivation may further impact eIF5A hypusination. Taken together, our results showed that eIF5A hypusination links polyamine pathway to autophagy activation and plays a central role in dynamic host-KSHV interactions regulating viral infection.

### eIF5A hypusination determines the translation efficiency of KSHV proteins

Hypusinated eIF5A plays a critical role in translation elongation by counteracting ribosome stalling during translation of difficult motifs, such as tri-proline (PPP) motif [67–69]. Schuller and colleagues recently described an extended repertoire of 29 motifs whose translation was dependent on hypusinated eIF5A (**Fig S6A**) [17]. The polyamine-hypusine axis is activated in multiple types of cancers, in which protein translation is hypusine-addicted to support cell growth and metastasis, while inhibitors targeting the polyamine-hypusine axis can be used for anti-cancer therapies [12, 14, 70, 71]. Since we also showed that KSHV infection modulates the polyamines-hypusine axis, we wondered whether eIF5A hypusination determines translation efficiency of KSHV proteins. We initially examined the protein sequence of KSHV ORF50/RTA (YP_001129401.1) for hyp-eIF5A-dependency motifs and identified 8 such motifs previously described by Schuller et al (**Fig 6A**). These sites are conserved across multiple KSHV strains (**Table S1**). We tested the impact of eIF5A hypusination on KSHV ORF50/RTA protein translation by using the DHPS-targeted eIF5A hypusination inhibitor, GC7 (N1-guanyl-1,7-diaminoheptane) [20, 72]. GC7 treatment led to the drastic reduction of Dox-induced KSHV RTA expression in iSLK.BAC16 cells (**Fig 6B**), which correlated with the reduction of hypusinated eIF5A as well as the inhibition of p62 accumulation, while GC7 caused neglectable cytotoxicity (**Fig S6B**). To further confirm, we ectopically expressed a cDNA of KSHV ORF50/RTA in SLK cells treated with GC7. Protein level of KSHV RTA was significantly lower due to GC7 treatment (**Fig 6C),**but its mRNA level was even moderately increased by GC7 (**Fig S6C**). On the contrary, we also tested GC7’s effect on expression of a GFP protein (CopGFP) that has no such difficult-to-translate motifs, which failed to lower protein level of GFP quantified by GFP-positive cells as well as MFI of GFP fluorescence (**Fig S6D,E**). Likewise, GC7 had no effect on EBV BZLF1/Zta protein level, since EBV BZLF1 only carries 2 hyp-eIF5A-dependency motifs (**Fig 6D, S6F**). Overall, these data confirmed that hypusinated eIF5A is important and specifically controls KSHV RTA protein translation.

**Figure 6.**
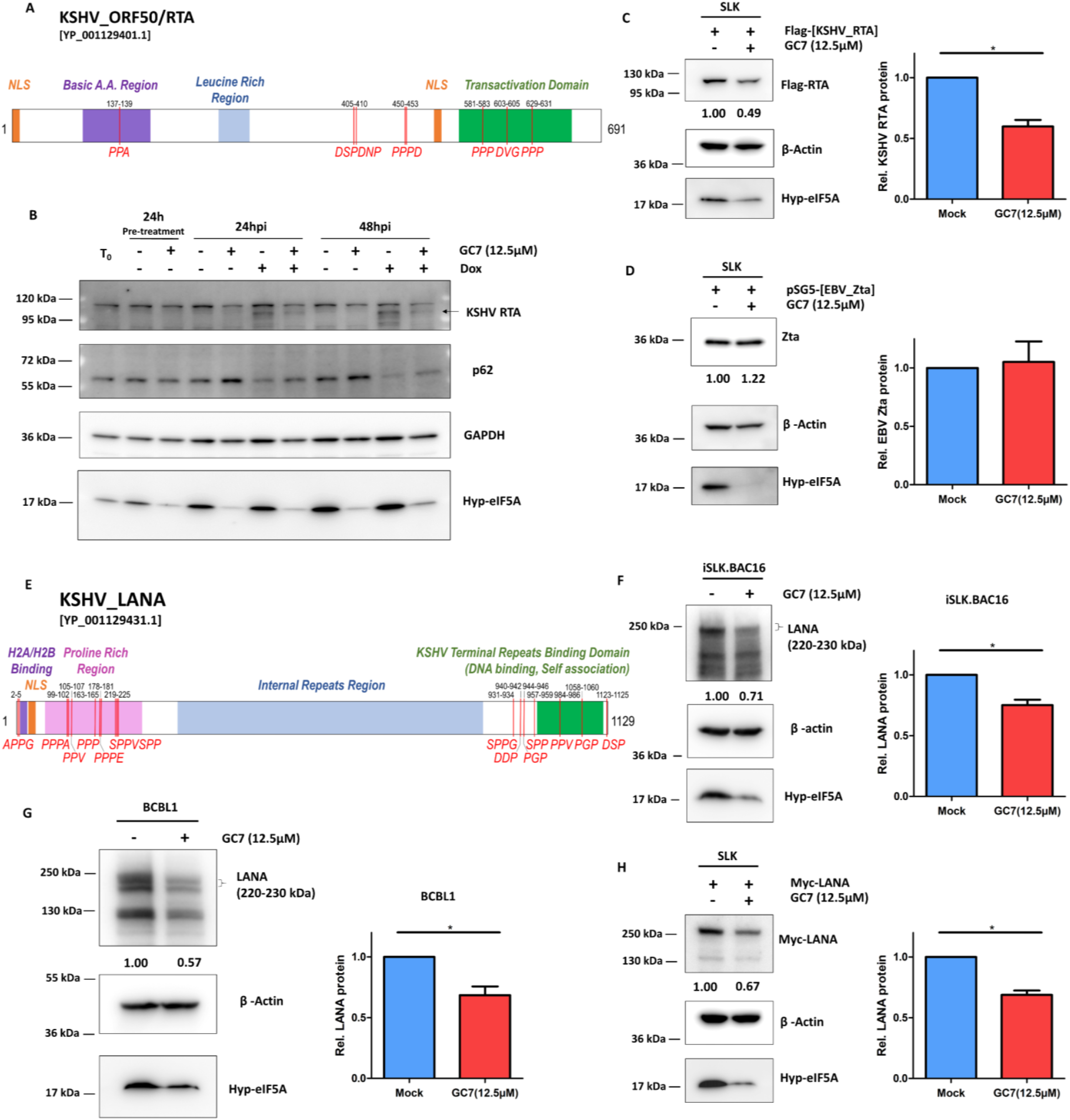
Translation of KSHV ORF50/RTA and LANA proteins requires eIF5A hypusination. (A). Schematic view of KSHV ORF50/RTA [YP_001129401.1] with hyp-eIF5A-dependent pause motifs annotated in red and several of its key domains. (B). Protein level of KSHV RTA, p62, and hyp-eIF5A in iSLK.BAC16 cells pre-treated with GC7 (12.5μM) for 24h and subsequently induced with Dox (1μg/mL) up to 48h was analyzed by immunoblotting assays. GAPDH was used as the loading control. (C, D). Protein level of KSHV RTA (C) or EBV Zta (D) along with hyp-eIF5A in SLK cells transfected respectively with the pcDNA vector expressing Flag-tagged KSHV RTA or non-tagged EBV Zta and subsequently treated with GC7 (12.5μM) for 48h was analyzed by immunoblotting assays. β-actin was used as the loading control. (E). Schematic view of KSHV ORF73/LANA [YP_001129431.1] with hyp-eIF5A-dependent pause motifs annotated in red and several of its key domains. (F, G). Protein level of KSHV LANA along with hyp-eIF5A in iSLK.BAC16 (F) or BCBL1 (G) cells treated with GC7 (12.5μM) for 48h was analyzed by immunoblotting assays. β-actin was used as the loading control. (H). Protein level of KSHV LANA along with hyp-eIF5A in SLK cells transfected with the pcDNA vector expressing myc-tagged KSHV LANA and subsequently treated with GC7 (12.5μM) for 48h was analyzed by immunoblotting assays. β-actin was used as a loading control. Intensity of KSHV protein bands was determined by using AlphaView SA (software) and normalized to β-actin. Results were calculated from on n=2-3 independent experiments and presented as mean ± SEM (* p<0.05; ** p<0.01; *** p<0.001, two-tailed paired Student t-test).

In parallel, we also examined the protein sequence of KSHV ORF73/LANA [YP_001129431.1] and identified 19 hyp-eIF5A-dependency motifs (**Fig 6E**), which are mostly conserved across multiple KSHV strains (**Table S2**). We tested the impact of GC7 on KSHV ORF73/LANA expression in KSHV latently infected cells, iSLK.BAC16 (**Fig 6F**) and BCBL1 (**Fig 6G**) cells. In both cases, GC7 led to the significant decrease of LANA protein. To further confirm, we ectopically expressed a cDNA of KSHV ORF73/LANA in SLK cells treated with GC7. Protein level of LANA was significantly lower due to GC7 treatment but not its mRNA level (**Fig 6H, S6C**). Therefore, our results showed that eIF5A hypusination is critical to KSHV infection since it determines the translation efficiency of KSHV key lytic and latent proteins (RTA, LANA).

### Inhibition of eIF5A hypusination potently restricts KSHV viral infection

Since we showed that translation of KSHV key lytic and latent proteins (RTA, LANA) is hypusine-addicted and can be efficiently blocked by GC7, we speculated that eIF5A hypusination would be a promising host target for developing anti-KSHV therapies. As a proof of principle, we tested the anti-KSHV potency of GC7 as an eIF5A hypusination inhibitor. Indeed, GC7 treatment suppressed the Dox-induced expression of KSHV lytic genes in a dose-dependent manner with similar IC_50_ = 10.0μM, 10.1μM, 7.5μM for ORF50/RTA, K8, ORF26 respectively (**Fig 7A**). It was confirmed by immunoblotting of KSHV ORF45 lytic protein in above cells (**Fig 7B**). Similarly, GC7 also suppressed KSHV lytic reactivation in Dox-treated TREx BCBL1-RTA cells in the dose-dependent manner (**Fig 7C,D**) without obvious cytotoxicity (**Fig S7A**). We also showed that GC7 severely impairs the amplification of KSHV viral genomes in these cells (**Fig 7E**). GC7 demonstrated comparable anti-KSHV effect in HEK293.r219 cells treated with TPA+NaB to induce KSHV reactivation (**Fig 7F, S7B**). However, GC7 treatment failed to inhibit EBV lytic gene expression in Akata/BX cells treated with human IgG to induce EBV reactivation (**Fig S7C**). Overall, these results demonstrated that inhibition of eIF5A hypusination by GC7 potently and specifically restricts KSHV lytic reactivation but not EBV.

**Figure 7.**
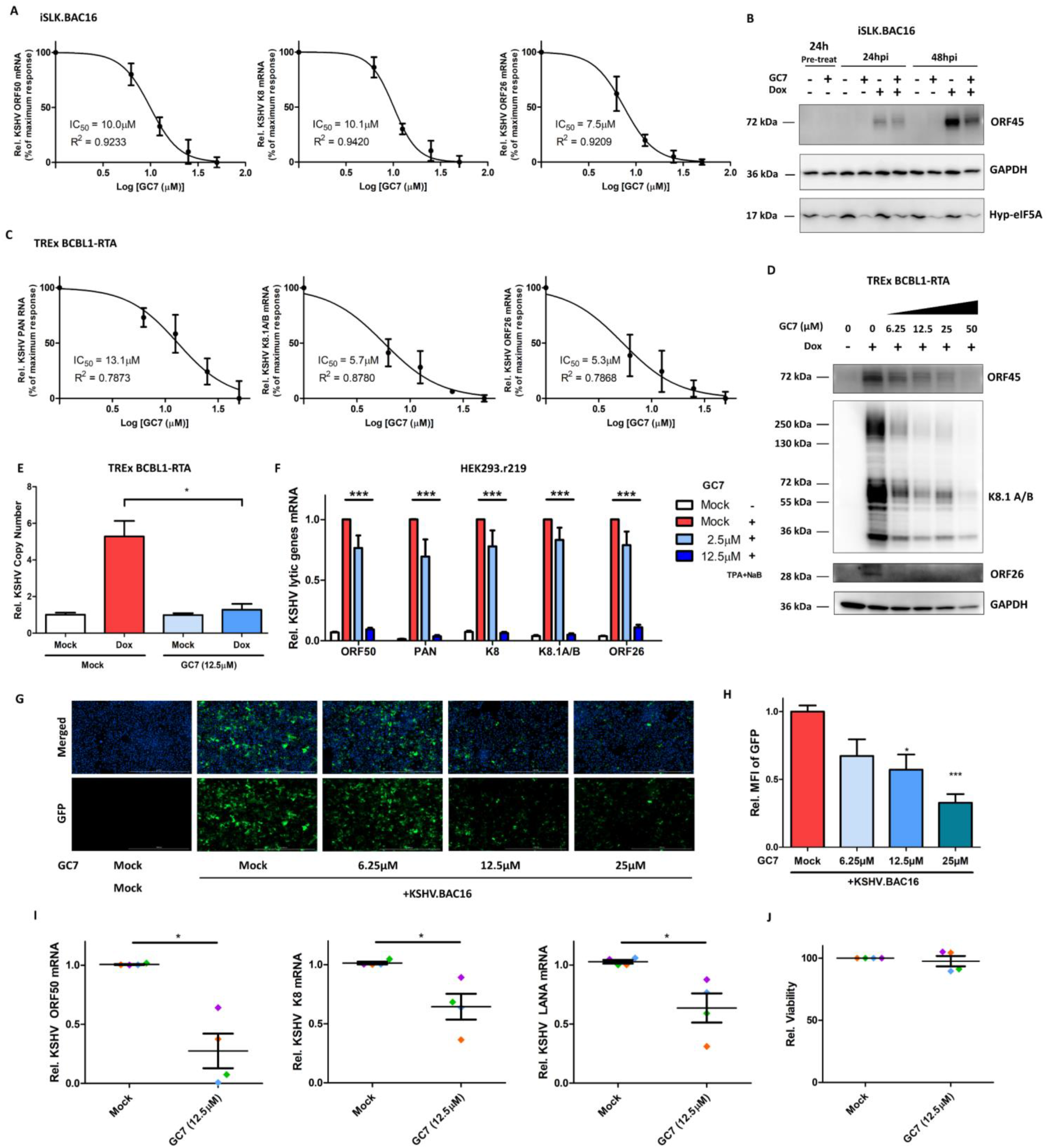
Inhibition of eIF5A hypusination efficiently blocks KSHV lytic infection. (A). mRNA level of KSHV lytic genes (ORF50/RTA, K8/K-bZIP and ORF26) in iSLK.BAC16 cells pre-treated with progressively increasing doses of GC7 for 24h and subsequently induced with Dox (1μg/mL) for 48h were analyzed by RT-qPCR assays and normalized to mock-treated induced cells. The drug effect at a series of doses was plotted as percentage of maximum response by using GraphPad PRISM 5. IC_50_ was determined for GC7’s inhibitory effect. (B). Protein level of KSHV ORF45 and hyp-eIF5A in iSLK.BAC16 cells pre-treated with GC7 (12.5μM) for 24h and subsequently induced with Dox (1μg/mL) up to 48h was analyzed by immunoblotting assays. GAPDH was used as the loading control. (C-E). mRNA level of KSHV lytic genes (PAN RNA, K8.1, ORF26) in TREx BCBL1-RTA cells pre-treated with progressively increasing doses of GC7 for 24h and subsequently induced with Dox (1μg/mL) for 48h were analyzed by RT-qPCR assays and normalized to mock-treated induced cells. The drug effect at a series of doses was plotted as percentage of maximum response by using GraphPad PRISM 5 (C). IC_50_ was determined for GC7’s inhibitory effect. Protein level of KSHV lytic genes (ORF45, K8.1A/B, ORF26) in above cells was analyzed by immunoblotting assays (D). GAPDH was used as the loading control. Relative copy number of KSHV viral genomes was also determined (E). (F). mRNA level of KSHV lytic genes (ORF50/RTA, PAN, K8, K8/K-bZIP, ORF26) in HEK293.r219 cells pre-treated with GC7 for 24h and subsequently induced with TPA (20 ng/mL) + NaB (0.3mM) for 48h was analyzed by RT-qPCR assays and normalized to mock-treated induced cells. (G, H). SLK cells were *de novo* infected with KSHV BAC16 viruses (MOI=0.5). Unbound viruses were washed away, and cells were subsequently treated with increasing doses of GC7 for 48h, then subjected to fluorescence imaging analysis (G). Nuclei were labelled with Hoechst (Blue), while GFP expression indicated infection with KSHV BAC16 viruses. MFI of GFP expression from five different fields of view for ≥7000 cells was measured and normalized to mock-treated cells (H). (I, J). Primary tonsillar B cells isolated from 4 healthy donors were *de novo* infected with KSHV BAC16 viruses (MOI=3). Unbound viruses were washed away, and cells were subsequently treated with GC7 (12.5 μM) for 72h, followed by RT-qPCR assays to measure mRNA level of KSHV lytic genes (ORF50/RTA, K8/K-bZIP) and latent gene (ORF73/LANA) and normalized to mock-treated cells (I). Cell viability was analyzed by CellTiter-Glo assays and normalized to mock-treated cells (J). Results were calculated from n=3 independent experiments and presented as mean ± SEM (* p<0.05; ** p<0.01; *** p<0.001, two-tailed paired Student t-test).

Furthermore, we determined the effect of GC7 on KSHV *de novo* infection. SLK cells were subjected to brief spin infection with KSHV.BAC16 viruses (MOI = 0.5). Unbound viruses were washed away, and cells were treated with GC7. KSHV infection rate was quantified by measuring the GFP expression from KSHV viral genomes in the infected cells. Fluorescence imaging showed that GC7 significantly reduces GFP expression in KSHV-infected SLK cells at 2 days post of infection (DPIs) in the dose-dependent manner (**Fig 7G,H**). We also performed the similar assays using primary tonsillar B cells, which have been shown susceptibility to KSHV infection and support active lytic replication of KSHV for up to 5 DPIs [49]. Hence, primary tonsillar B cells from healthy donors were isolated and subjected to brief spin infection with KSHV.BAC16 viruses (MOI = 3). Unbound viruses were washed away, and cells were treated with GC7. Expression of KSHV lytic (ORF50/RTA, K8) and latent (ORF73/LANA) genes at 3DPIs were analyzed by qPCR assays, which clearly showed that GC7 significantly reduces KSHV viral gene expression in tonsillar B cells from four donors (**Fig 7I**) without any cytotoxicity (**Fig 7J**). These results suggested that eIF5A hypusination also participates in the early steps of viral life cycle during KSHV *de novo* infection and that its inhibition by GC7 potently blocks early events and prevents establishment of following persistent KSHV infection.

## Discussion

Our innovative studies demonstrated that polyamine synthesis and eIF5A hypusination are dynamically regulated by KSHV infection. We observed that intracellular polyamines are overall upregulated in KSHV latently infected tumor cells by PolyamineRED staining (**Fig 1**). However, the results from TLC assays have certain discrepancy likely due to that this type of analysis only accesses to free polyamines and that disruption of cells may cause the significant loss of polyamines as well. Nevertheless, TLC assays showed that spermine, the polyamine species with the highest molecular weight, is still more abundant in KSHV latently infected tumor cells. Impact of KSHV latency on polyamine synthesis could be explained by the fact that the rate-limiting enzyme ODC1 is upregulated (**Fig 1**), since ODC1 is subjected to transcriptional regulation of c-Myc [45, 46] that is upregulated by KSHV ORF73/LANA protein dominantly expressed in KSHV latency [44]. Interestingly, spermidine is somehow lowered in KSHV latently infected cells in TLC assays, which could be explained by spermidine being consumed for the generation of hypusinated eIF5A that is indeed present at higher level in KSHV latently infected cells (**Fig 5**). This may benefit the translation of KSHV key latent protein ORF73/LANA, which is shown to depend on eIF5A hypusination in our own studies (**Fig 6**). On the contrary, KSHV lytic reactivation triggers an overall decrease of intracellular polyamines, particularly a dramatic depletion of spermidine (**Fig 1**), correlating with the decrease of ODC1 during KSHV lytic reactivation (**Fig 1**) as well as the further increase of hypusinated eIF5A in KSHV-reactivated cells (**Fig S5**). This is likely required to fulfill the even higher demand of KSHV viral protein translation during lytic reactivation, since the majority of KSHV genes (over 100) is expressed in lytic phase while only a dozen in latent phase. We indeed showed that eIF5A hypusination is critical to translation of KSHV key lytic protein ORF50/RTA (**Fig 6**). Other KSHV lytic proteins may also depend on hypusinated eIF5A to translate, which needs further investigation. There could be also other explanations for KSHV reactivation induced spermidine drop. For instance, polyamines are strongly charged polycations, and it has been reported that spermidine and spermine are packed in HSV-1 virions [21], allegedly to neutralize the negative charge of its large viral genomes and help with DNA compaction. It still remains uncharacterized whether spermidine is also incorporated into KSHV virion as HSV-1.

In these studies, we also clearly showed that polyamine synthesis and eIF5A hypusination are critically required for KSHV infection. RNAi-mediated knockdown of several key enzymes participating in polyamine synthesis, including ODC1, AGMAT, SRM, and SMS, unanimously impairs KSHV lytic reactivation in multiple KSHV-infected cell systems (**Fig 2,3**), which is further supported by the results that several chemical probes inhibiting polyamine synthesis, including DFMO, clofazimine (CLF), and ribavirin, all block KSHV lytic reactivation in these cells (**Fig 3,4**). Furthermore, our studies also innovatively identified that eIF5A hypusination is the key linker to connect polyamine pathway with autophagy activation that is also critical to KSHV viral infection, as we observed that ODC1 inhibitor DFMO indeed represses eIF5A hypusination as well as autophagy activation that favors KSHV lytic switch (**Fig 5**). In addition, a supportive finding is that knockdown of ODC1 seems to preferentially decrease spermidine that is the key polyamine species utilized for eIF5A hypusination (**Fig 2**). Likewise, we demonstrated that RNAi-mediated knockdown of the key enzyme DHPS participating in eIF5A hypusination as well as its inhibition by GC7 both efficiently suppress KSHV lytic reactivation in multiple KSHV-infected cell systems (**Fig 5, 7**). We further unraveled that one potential mechanism is that translation of certain KSHV viral proteins could be hypusine-addicted (**Fig 6**), as activation of eIF5A depends on its hypusination by using spermidine [16], which enables eIF5A to alleviate ribosome stalling on defined hard-to-translate tri-peptide motifs [17, 18]. We indeed recognized that KSHV ORF50/RTA and ORF73/LANA proteins carry multiple of such motifs that are conserved across all KSHV strains (**Fig 6, Table S1,2**). Beyond KSHV lytic reactivation, inhibition of eIF5A hypusination by GC7 also impairs KSHV *de novo* infection, likely through affecting synthesis of certain KSHV viral proteins possessing immune antagonizing or cell proliferating functions which are required for early steps of KSHV infection. For example, NF-κB is activated shortly post of KSHV *de novo* infection and important to the establishment of latency [73].

By combining our new results with earlier knowledge regarding to host and viral controls of KSHV infection, we propose the following model describing the comprehensive roles of polyamine pathway and eIF5A hypusination in regulating KSHV viral infection (**Fig S8**). At the latent phase of KSHV infection, LANA upregulates ODC1 expression while suppresses RTA transcription [74]. ODC1 upregulation promotes intracellular polyamine biosynthesis leading to an increase of spermine. In contrast, spermidine is actively and steadily consumed, and hypusinated eIF5A is accumulated to further promote translation of LANA. At the early stage of lytic switch, RTA transcription is activated, and the newly transcribed RTA mRNAs can thus be efficiently translated thanks to the pre-existing higher level of hypusinated eIF5A. RTA in turn promotes autophagy activation that would also benefit from increased hypusinated eIF5A [19, 20], which overall favors KSHV lytic switch. However, at the late stage of lytic phase, RTA expression is strong enough to counteract LANA, which leads to the reduction of ODC1. Additionally, it has been shown that c-Myc expression is transcriptionally downregulated by the KSHV lytic gene vIRF4 [48], which would also contribute to the decrease of ODC1. ODC1 reduction decreases intracellular polyamines, and thus hamper cell cycle progression [75], which has been shown to favor KSHV lytic reactivation [76]. In addition, accumulated spermine would convert to spermidine and putrescine through SMOX and PAOX, which further generates hydrogen peroxide and oxidative stress that have also been shown to promote KSHV lytic reactivation [77, 78]. At this moment, KSHV lytic protein synthesis, especially RTA, is already less dependent on hypusinated eIF5A, so decrease of intracellular polyamines may generate much less adverse effect on KSHV lytic protein synthesis. Additionally, the resulting attenuated autophagy would actually benefit the late phase of KSHV lytic replication [79].

**Figure 8.**
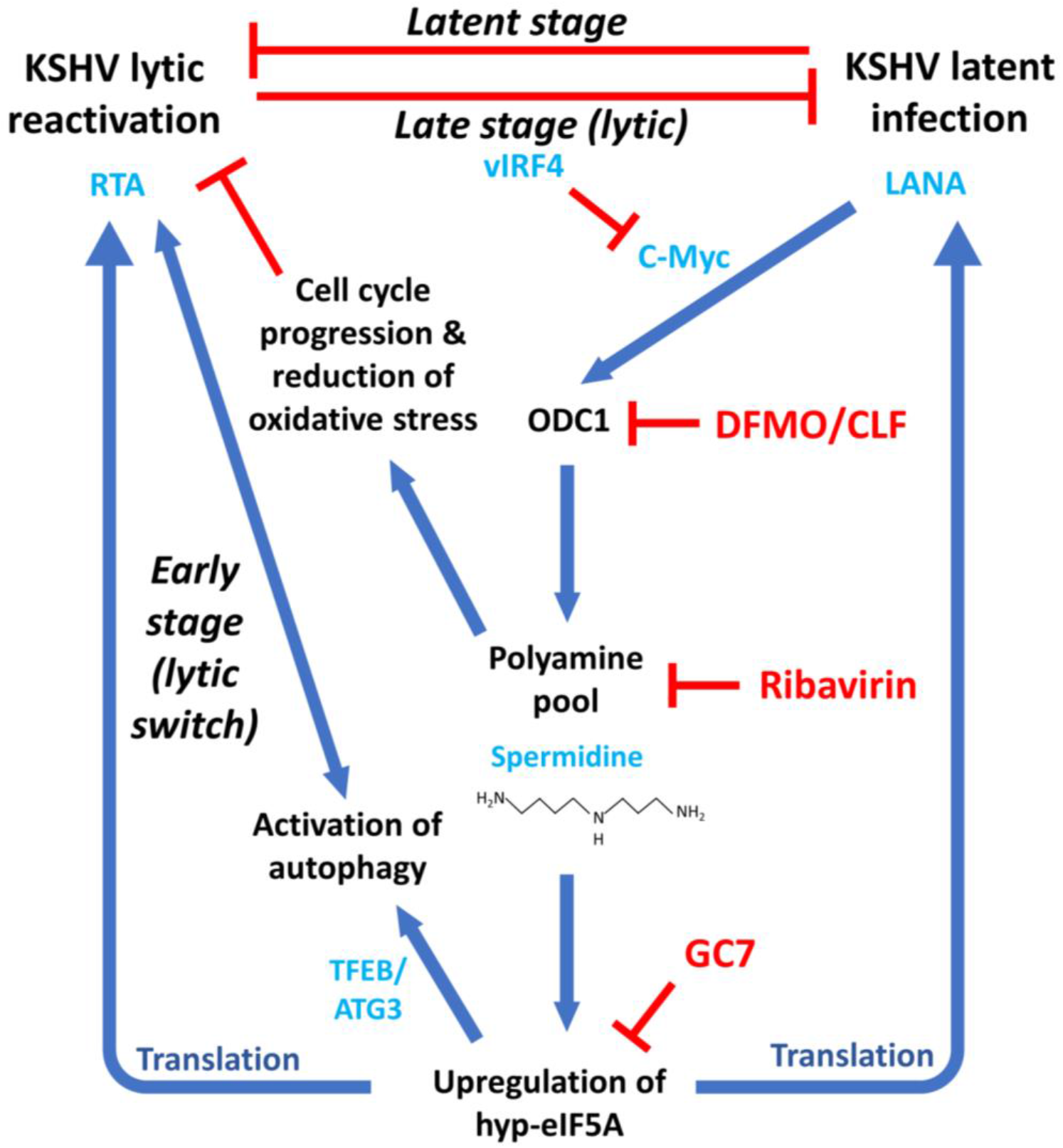
A proposed model illustrates the dynamic and profound interaction of host polyamine biosynthesis and eIF5A hypusination with KSHV infection. At the stage of latent infection, KSHV ORF73/LANA protein induces upregulation of c-Myc that further transcriptionally activates ODC1, which results in the overall increase of intracellular polyamines. However, spermidine is consumed to produce hypusine-eIF5A ensuring the efficient translation of LANA protein required for maintenance of KSHV latency. This results in a positive feedback sustaining activation of the polyamine-hypusine axis. At the early stage of lytic switch, KSHV ORF50/RTA gene starts to be actively transcribed upon certain stimuli, and the constant high level of hypusine-eIF5A not only ensures the efficient translation of RTA protein but also facilitates the activation of autophagy, which overall promotes KSHV lytic reactivation. On the contrary, at the late stage of lytic replication KSHV RTA protein is expressed up to a sufficient level to suppress LANA protein, leading to decrease of ODC1 and reduction of intracellular polyamines. In addition, other KSHV lytic proteins, such as vIRF4, also contribute to the downregulation of c-Myc, causing further decrease of ODC1 and polyamines. This would generate an adverse effect on cell cycle progression and an increase of oxidative stress, which would be beneficial to KSHV lytic reactivation. Thus, cells are now adjusted to a favorable environment to further promote KSHV lytic replication and viral propagation. KSHV *de novo* infection likely suppresses ODC1 as well to allow early steps of viral replication. Given to the critical roles of polyamine biosynthesis and eIF5A hypusination in KSHV infection, inhibitors targeting these metabolic pathways would serve as novel antiviral reagents to efficiently block both KSHV latent and lytic replications.

The evidence is emerging that KSHV modulates cellular metabolic landscape in order to fulfill its particular needs to promote viral propagation and oncogenesis [37, 38]. A recent study [39] reported that broad modifications of host metabolism, including induction of arginine and proline metabolism, occur in KSHV-infected cells in 3D culture. Arginine is a precursor metabolite for polyamine synthesis (**Fig 2A**), and it can be sequentially converted to putrescine via ODC1 and AGAMT, both of which were identified as critical to KSHV lytic reactivation in our studies (**Fig 2, 3**). Furthermore, these authors also reported that ornithine, the direct substrate of ODC1, is induced in KSHV-infected cells, which resonates with our finding that polyamines are upregulated by KSHV latent infection. Overall, these observations along with ours illustrated that KSHV profoundly reprograms the metabolic pathways of host cells to not only promote viral latency and oncogenesis but also favor viral lytic reactivation and dissemination of KSHV viruses. However, it is still not fully clear how KSHV infection modulates polyamine pathway and eIF5A hypusination. We believe that regulation of ODC1 by KSHV LANA might be just one of mechanisms. Some large DNA viruses encode functional genes related to the polyamine metabolism, such as *Paramecium bursaria* chlorella virus (PBCV-1) virus that encodes genes equivalent to most of the biosynthesis pathway [80–82] in its 331-kb long genome. More recently, it has been reported that the viral genome of bovine gammaherpesvirus BoVH-6 encodes a viral protein similar to ODC1 [83]. There are nearly 100 KSHV-encoded proteins, some of which may directly impact on polyamine pathway and eIF5A hypusination. Such functions of certain KSHV proteins are pending for discovery.

After all, our understanding of polyamine pathway and eIF5A hypusination in the context of KSHV infection will pave the way for development of new antiviral and antitumor strategies to treat KSHV-associated malignancy. As the proof of principle in our studies, inhibitors targeting polyamine pathway (DFMO, clofazimine, ribavirin) and eIF5A hypusination (GC7) can be used as antivirals to block latent and lytic replications as well as *de novo* infection of KSHV (**Fig S8**). As the matter of fact, three of them are already FDA-approved drugs currently being prescribed for treating other diseases. Thus, these drugs are safe for future evaluation of their potential to treat KSHV-associated malignancy in clinic. Beyond inhibition of KSHV latent infection, interruption of KSHV lytic infection is equally important to not only block generation of new KSHV virions and their dissemination but also reduce viral tumorigenesis since KSHV lytic infection contributes notably to tumors as well by generating angiogenesis and anti-apoptotic phenotypes (reviewed in [40, 41]). In conclusion, polyamine pathway and eIF5A hypusination play a comprehensive role in regulating KSHV infections, which serve as promising new drug targets for treating KSHV infection and KSHV-associated malignancy.

## Materials and Methods

### Cell culture

HEK293.r219 [50] and iSLK.BAC16 [42] cells were respectively obtained from Dr. Prashant Desai (Johns Hopkins) and Dr. Shou-Jiang Gao (University of Pittsburgh) and cultured in DMEM supplemented with 10% FBS under selection (HEK293.r219: 5 μg/mL puromycin; iSLK.BAC16: 1.2 mg/mL Hygromycin B, 2 μg/mL puromycin, 250 μg/mL G418). TREx BCBL1-RTA [84] cells were obtained from Dr. Jae Jung (Cleveland Clinic) and maintained in RPMI supplemented with 10% FBS under selection (200 μg/mL Hygromycin B). BC-3, BCBL1 and SLK cells were acquired from the NIH AIDS Reagent Program. These cells were maintained in RPMI supplemented with 10% FBS (BC-3 with 20% FBS). BJAB and Akata/BX cells were acquired from Dr. Renfeng Li (Virginia Commonwealth University) and maintained in RPMI supplemented with 10% FB S (Akata/BX under selection 500μg/mL G418). Tonsillar mononuclear cells and B lymphocytes were maintained in RPMI (Gibco) supplemented with 10% heat-inactivated human serum (Advanced Biotechnologies Inc, Cat#P2-203), 100 U/mL penicillin, 100 μg/mL streptomycin, 0.5 μg/mL Amphotericin B, and 50 μM β-Mercaptoethanol as previously described [85]. KSHV lytic reactivation was induced in TREx BCBL1-RTA cells with 1μg/mL doxycycline, in iSLK.BAC16 cells with either 1μg/mL doxycycline alone or in combination with 1mM Sodium Butyrate (NaB), and in HEK293.r219 cells with 20ng/mL 12-O-Tetradecanoylphorbol 13-acetate (TPA) and 0.3mM NaB or ectopic expression of KSHV ORF50/RTA cDNA.

### Compounds

12-O-Tetradecanoylphorbol 13-acetate (Cat#P8139) and sodium butyrate (Cat#AAA1107906) were purchased from Sigma-Aldrich and Fisher Scientific, respectively. Doxycycline was obtained from Fisher Scientific (Cat#BP2653-1). Polyamine supplement (1000x) (Cat#P8483), putrescine (Cat#P5780), spermidine (Cat#S0266), spermine (Cat#S4264), and clofazimine (Cat# C8895) were purchased from Sigma-Aldrich. 2-difluoromethylornithine (DFMO) (Cat#2761), ribavirin (Cat#R0077), and deoxyhypusine synthase inhibitor N1-guanyl-1,7-diaminoheptane (GC7) (Cat#259545) were purchased from Tocris, Tokyo Chemical Industry, and EMD Millipore respectively.

### Isolation of tonsillar B lymphocytes

Tonsillar tissues were acquired through the National Disease Research Interchange (NDRI, Philadelphia) and were collected from de-identified healthy donors via routine tonsillectomy procedures. Extraction of mononuclear cells from the tonsillar tissue was performed as previously described [85]. In brief, the block of tissue was cut into smaller (3mm) pieces, and cells were mechanically dissociated by using a stainless-steel sieve (250-μm mesh) with a syringe plunger, through which the tissue samples remained cold and moistened in the Hanks balanced salt solution (HBSS) with antibiotics (100 U/mL penicillin, 100 μg/mL streptomycin, 5 μg/mL gentamicin, 0.5 μg/mL Amphotericin B). Cell suspension was passed through a 40 μm plastic cell strainer and overlaid onto 10 mL Ficoll-Hypaque (GE Healthcare), followed by centrifugation (1000x g) for 20 mins at 4°C. Mononuclear cells were collected from the buffy coat at the interface, and washed three times with HBSS containing antibiotics and centrifuged (300x g) for 10mins at 4°C. Washed cells were resuspended in cryopreservation media (HyClone, Cat#SR30001.02) mixed with tonsil mononuclear cells complete media without antibiotics. B lymphocyte isolation was conducted through negative selection by using the B cell Isolation Kit II (Miltenyi Biotec, Cat#130-091-151) according to the manufacturer’s protocol. Purity of B lymphocytes (>98%) was assessed via immunostaining by using an anti-human-CD19, PE-conjugated antibody (Miltenyi Biotec, Cat#130-113-731) per manufacturer’s instructions and analyzed via flow cytometry on an Accuri C6 Plus (BD Biosciences).

### Preparation of KSHV viruses and de novo infection

iSLK.BAC16 cells were treated with 1μg/mL doxycycline and 1mM NaB for 24h, and then kept in fresh media only containing 1μg/mL doxycycline. Doxycycline was refreshed every 2 days. Supernatants were collected 6 dpi, centrifuged (400x g) for 10 mins to remove cellular debris, filtered through the 0.45μm filter and stored at −80°C. KSHV BAC16 viruses were titrated in HEK293T cells through the serial dilution of viral stock along with 8μg/mL polybrene via spinoculation (2500 rpm) for ~2h at 37°C. Percentage of GFP-positive cells was determined by flow cytometry on an Accuri C6 Plus. Forward Scatter (FS) and Side Scatter (SS) were used for gating of single cells. Analysis was performed using the FlowJo v10 software. Primary tonsillar B lymphocytes were spinoculated (2500 rpm) with KSHV BAC16 viruses (MOI=3) along with 8μg/mL polybrene for ~2h at 37°C. Cell media was completely removed at 12hpi. Unbound viruses were washed away, and fresh media containing the tested compounds were added. Similarly, SLK cells were subjected to KSHV *de novo* infection (MOI=0.5).

### Thin-layer chromatography

Protocol of thin-layer chromatography (TLC) to analyze polyamines was adapted from previous report [24, 86, 87]. Cells were harvested and resuspended in 2% perchloric acid solution (v/v) in ddH2O (Cat#SP339-500), sonicated at 4°C (30% amplitude, 1min total with 2sec ON and 2sec OFF), and incubated at 4°C for overnight on a rotating shaker. Cell lysates were centrifuged (11500x g) for 45mins at 4°C, and transferred to a new test tube. The following solutions were prepared freshly. 1 volume of cell lysates was mixed with 2 volumes of dansyl-chloride solution (18.6mM in Acetone; Cat#D-2625, Cat#BP392-100)) and 1 volume of supersaturated Sodium Carbonate solution (4.44M in ddH2O; Cat#S263-500), which was vortexed thoroughly and incubated for overnight in dark on a rotating shaker at room temperature. 0.5 volume of L-proline solution (1.3M in ddH2O; Cat#BP392-100) was added, then samples were thoroughly vortexed and incubated for another 1h. For extraction of dansylated polyamines, 2.5 volume of toluene (Cat# AC610951000) was added, and the mixtures were thoroughly vortexed and further incubated for 20 mins. The mixtures were centrifuged (13500 rpm) for 10mins at room temperature. The organic phase (upper layer) was carefully separated and spotted (~4uL) onto a TLC plate (Cat#1.05721.0001), and individual polyamines were separated by ascending chromatography using a cyclohexane:ethylacetate mixture (2:3) (v:v) as the eluant for 30 mins. Standards and controls were processed identically in parallel. The TLC plate was then imaged by using a FluorChem E (ProteinSimple) under UV light exposure.

### Cell transfection

Reverse transfection of siRNA was performed using Lipofectamine RNAiMAX (Invitrogen) as previously described [88]. Following Silencer Select siRNAs (Invitrogen) were used: ornithine decarboxylase 1 (ODC1) (s9821, s9822, s9823 as si1-3), agmatinase (AGMAT) (s379, s381 as si1-2), spermidine synthase (SMS) (s13173, s13175 as si1-2), spermine synthase (SRM) (s13430, s13432 as si1-2), deoxyhypusine synthase (DHPS) (s91, s92 as si1-2), and non-targeting control (Cat#AM4641). Turbofect (ThermoFisher) was used for vector transfection following manufacturer’s recommendations. Flag-tagged KSHV ORF50/RTA vector was a gift from Dr. Pinghui Feng (University of South California) [89], pSG5-BZLF1 was a gift from Dr. Diane Hayward (Addgene # 72637) [90], and pA3M-LANA was a gift from Dr. Erle Robertson (University of Pennsylvania). The pmaxGFP plasmid was obtained from Lonza.

### Immunoblotting

Protein immunoblotting was performed as previously described [91]. The following antibodies were used: anti-ODC1 (EMD Millipore, Cat#MABS36), anti-hypusine-eIF5A (EMD Millipore, Cat#ABS1064), anti-SSAT-1 (Novus, Cat#NB110-41622), anti-LC3I/II (CST, Cat#4108), anti-SQSTM1/p62 (D5E2) (CST, Cat#8025), anti-GAPDH (SCBT, Cat#sc-47724), anti-β-actin (SCBT, Cat#sc-47778), anti-DHPS (SCBT, Cat#sc-365077), anti-EBV-ZEBRA/Zta (SCBT, Cat#sc-53904), anti-HHV8-K8.1A/B (SCBT, Cat#sc-65446), anti-HHV8-ORF50/RTA (Abbiotec, Cat#251345), anti-KSHV-ORF26 (2F6B8) (Novus, Cat#NBP1-47357), anti-KSHV-LANA (Advanced Biotechnologies, cat# 13-210-100), antiFLAG (CST, Cat#14793), anti-mouse-HRP (Invitrogen, Cat# 31430), anti-rabbit-HRP (Invitrogen, Cat#A27036), anti-rat-HRP (Invitrogen, Cat#A18739). An anti-KSHV-ORF45 antibody was a gift from Dr. Fanxiu Zhu (Florida State University).

### Real-Time quantitative PCR

The procedures of RNA extraction, cDNA preparation, genomic DNA extraction, and real-time quantitative PCR (RT-qPCR) assays were conducted as previously described [88]. Primers used in this study are listed in **Table S3**.

### Cell viability

CellTiter-Glo (Promega, Cat# G7570) was used for cell viability assays following the manufacturer’s instructions. Luminescence was measured on a BioTeK Cytation 5 plate reader.

### Confocal and fluorescence microscopy

SLK and iSLK.BAC16 cells were seeded onto the high precision cover glasses (Bioscience Tools, Cat#CSHP-No1.5-13) pre-coated with poly-L-lysine (R&D Systems, Cat#3438-100-01) for 45mins in a 24-well plate. These cells were induced with 1μg/mL doxycycline for 48h, and incubated with fresh media containing 10mM Polyamine Red (Funakoshi, Cat#FDV-0020) for 15mins. The following steps were all performed in dark. Cells were rinsed and fixed with 4% paraformaldehyde for 8 mins at room temperature, followed by staining with Hoechst (Invitrogen) per manufacturer’s guidelines.

Coverslips were rinsed and mounted on slides by using ProLong Glass Antifade Mountant (Invitrogen, Cat#P36982). Slides were left to cure in dark for 24h at room temperature per manufacturer’s recommendations. Confocal images were acquired by using the ZEISS LSM 700 Upright laser scanning confocal microscope and ZEN imaging software (ZEISS). Polyamines were imaged via TAMRA channel [43], while GFP is constitutively expressed in cells infected with KSHV BAC16 [42]. Image analysis was performed by using the ImageJ software. For BJAB and BCBL-1 cells, the procedure was similar with slight modifications: The suspended cells (4 × 10^4^ cells/mL) were incubated with 10mM PolyamineRED (Funakoshi Co.,Ltd) for 15 mins, and washed 2 times with 1x PBS. Cells were fixed and subjected to Hoechst staining, followed by the PBS washing. These cells were spun down onto the poly-L-lysine-coated high precision cover glasses in 24-well plates via brief centrifugation (2000 rpm) for 2mins. Cover glasses were mounted and analyzed as above. Fluorescence images of HEK293.r219 cells at GFP and RFP channels as well as SLK cells *de novo* infected with KSHV BAC16 viruses at GFP channel were acquired by using a BioTeK Cytation 5 plate reader. Hoechst was used to stain nuclei.

### Protein sequence scanning

Protein sequences were retrieved from the NCBI database and scanned for hyp-eIF5A-dependent pause motifs [17] by using the ScanProsite webtool (https://prosite.expasy.org/prosite.html) [92].

### Statistical analysis

Statistical analysis was performed by using GraphPad PRISM 5 or Excel. Results were presented as mean ± SEM. p-values were determined by using the twotailed paired Student t-test.

## Acknowledgments

We would like to thank Drs Prashant Desai, Jae Jung, Shou-Jiang Gao, Renfeng Li for the gifts of HEK293.r219, TREx BCBL1-RTA, iSLK.BAC16, and Akata/BX cells. SLK and BCBL-1 cells were acquired through the NIH AIDS Reagent Program thanks to Drs. Jay A. Levy and Sophie Leventon-Kriss for SLK, and Drs. Michael McGrath and Don Ganem for BCBL-1. We would like to thank Drs Pinghui Feng, Erle Robertson, and Diane Hayward for the gifts of Flag-tagged KSHV-ORF50/RTA, pA3M-LANA, and pSG5-BZLF1 vectors. We would also like to thank Dr. Fanxiu Zhu for the gift of anti-KSHV-ORF45 antibody.

## Author contributions

J.Z., G.F. conceived the study. G.F. performed experiments. G.F., J.Z., N.S. analyzed data and wrote the manuscript. A.B., D.Z., W.K. and M.J. and provided reagents, technical support, or comments of this study. J.Z. provided overall supervision of the study.

## Data Availability statement

The authors declare that all relevant data supporting the study can be found within the manuscript or the supplementary information.

---

**Figure S1.**
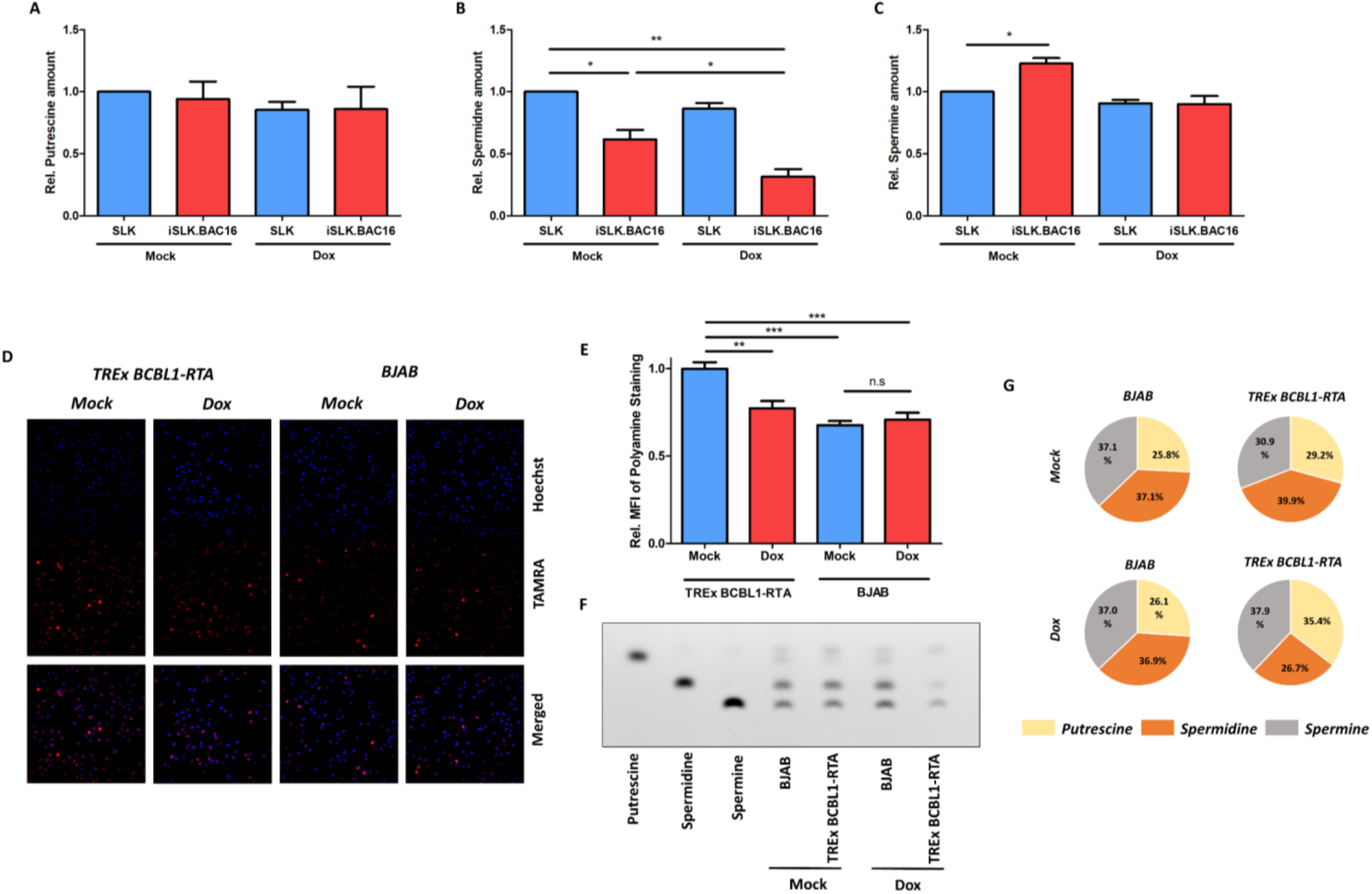
(A-C). Relative abundance of putrescine (A), spermidine (B), and spermine (C) in Figure 1C. (D, E). Intracellular polyamines in TREx BCBL1-RTA and BJAB cells treated with Dox for 48h or mock were stained with PolyamineRED (TAMRA) and visualized by confocal fluorescence microscopy (D), whereas nuclei were labelled with Hoechst (Blue). Mean fluorescence intensity (MFI) of polyamine fluorescence signal (TAMRA) of ≥ 2400 cells (E) was determined and normalized to mock-treated TREx BCBL1-RTA cells. (F, G). Intracellular polyamine species (putrescine, spermidine, spermine) in TREx BCBL1-RTA and BJAB cells treated with Dox for 48h or mock were analyzed by thin-layer chromatography (TLC) with pure individual polyamine species as reference (F). Relative changes of polyamine species in TLC results were displayed as the relative proportion of total intracellular polyamines in pie chart (G). Results were calculated from n=2-3 independent experiments and presented as mean ± SEM (* p<0.05; ** p<0.01; two-tailed paired Student t-test).

**Figure S2.**
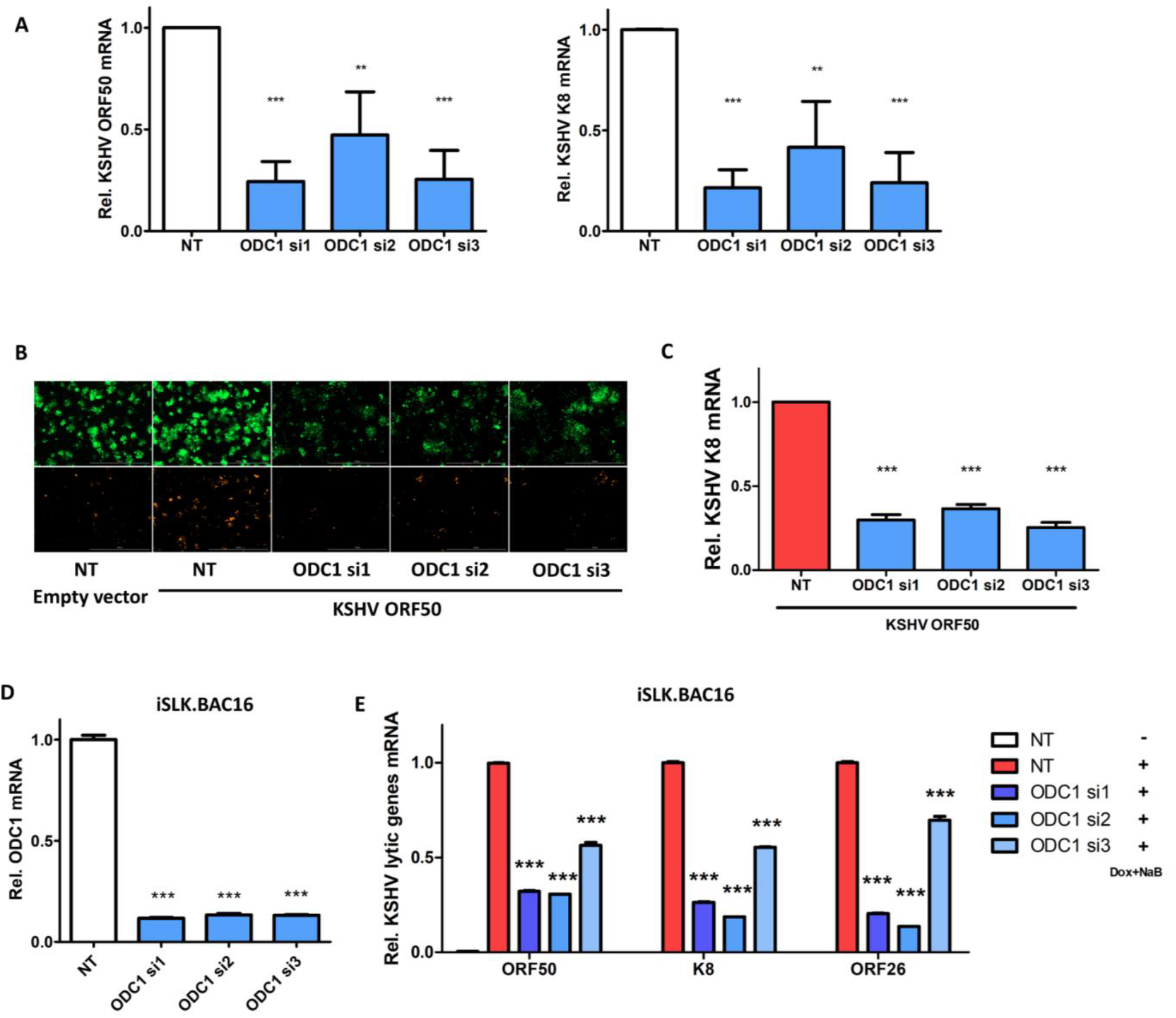
(A). mRNA level of KSHV lytic genes (ORF50/RTA, K8) in un-induced HEK293.r219 cells transfected with ODC1 siRNAs (si1, si2, si3) or non-targeting control siRNA (NT) was analyzed by RT-qPCR assays and normalized to NT-transfected cells. (B, C). HEK293.r219 cells transfected with ODC1 siRNAs (si1, si2, si3) or NT were induced with ectopic expression of ORF50 for 48h and visualized by fluorescence imaging (B). mRNA level of KSHV lytic gene K8 in above cells was analyzed by RT-qPCR assays and normalized to NT-transfected ORF50-induced cells. (D, E). mRNA level of ODC1 (D) and KSHV lytic genes (ORF50/RTA, K8, ORF26; E) in iSLK.BAC16 cells transfected with ODC1 siRNAs (si1, si2, si3) or NT and subsequently induced with Dox (1μg/mL) + NaB (1mM) for 48h was analyzed by RT-qPCR assays and normalized to NT-transfected induced cells. Results were calculated from n=3 independent experiments and presented as mean ± SEM (* p<0.05; ** p<0.01; *** p<0.001; two-tailed paired Student t-test).

**Figure S3.**
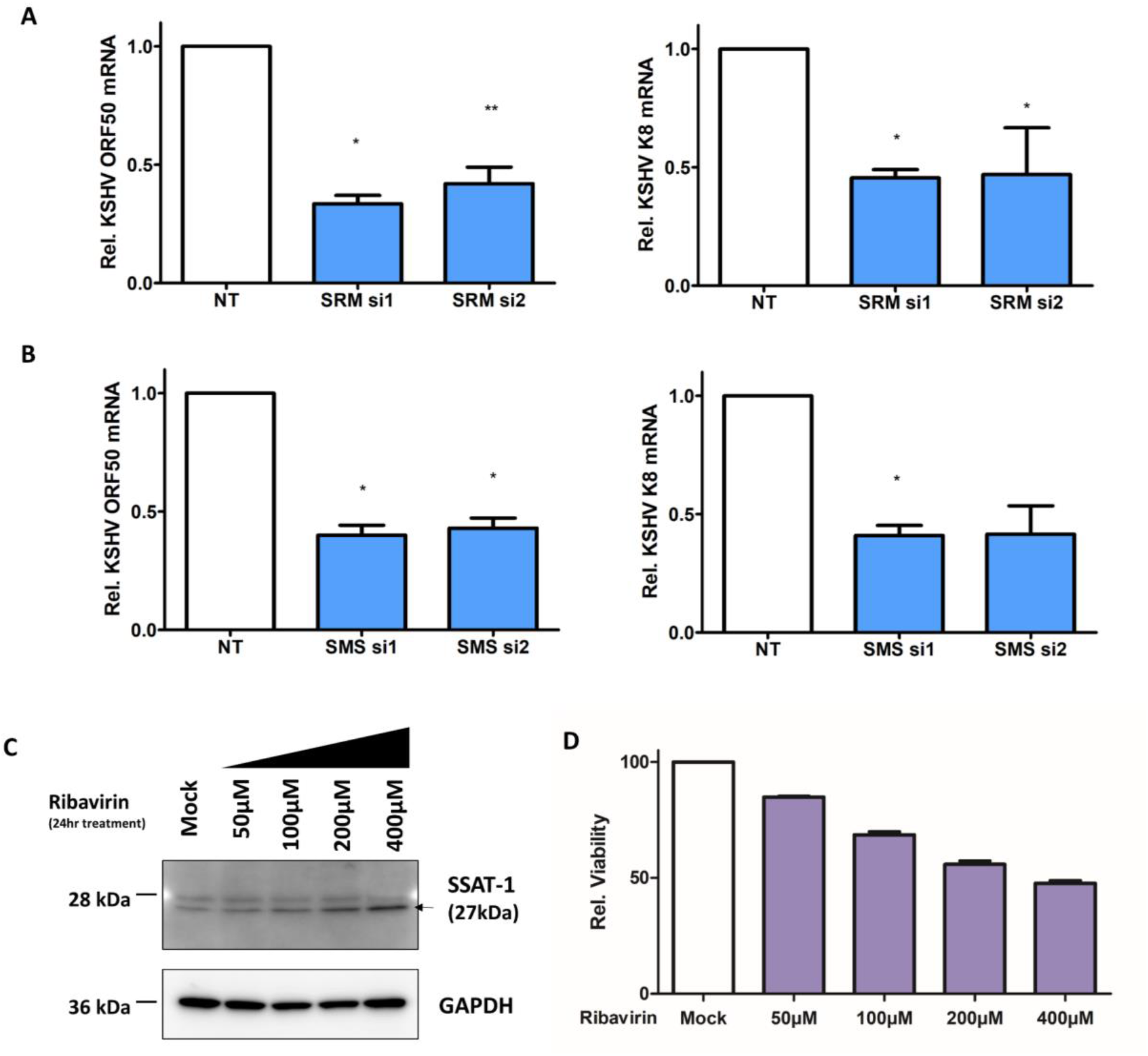
(A, B). mRNA level of KSHV lytic genes (ORF50/RTA, K8) in un-induced HEK293.r219 cells transfected with siRNAs targeting SRM (si1, si2; A) or SMS (si1, si2; B), or NT was analyzed by qRT-assays and normalized to NT-transfected cells. (C, D). Protein level of SSAT-1 in iSLK.BAC16 cells treated with increasing doses of ribavirin for 24h was analyzed by immunoblotting assays (C). GAPDH was used as the loading control. Cell viability of above cells was analyzed by CellTiter-Glo assays and normalized to mock-treated cells (D). Results were calculated from n=2-3 independent experiments and presented as mean ± SEM (* p<0.05; ** p<0.01; two-tailed paired Student t-test).

**Figure S4.**
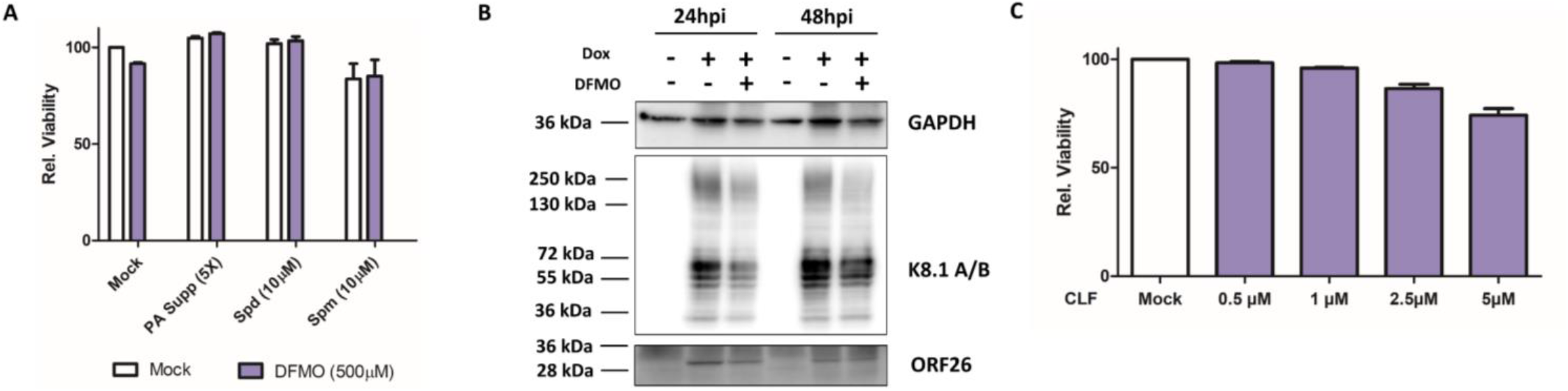
(A). Cell viability of HEK293.r219 cells treated with or without exogenous polyamines (mixed polyamines supplement [PA Supp, 5x], Spermidine [Spd, 10μM] or Spermine [Spm, 10μM]) in the presence or absence of DFMO (500μM) for 24h was analyzed by CellTiter-Glo assays and normalized to mock-treated cells. (B). Protein level of KSHV lytic genes (K8.1A/B, ORF26) in TREx BCBL1-RTA cells pre-treated with DFMO (500μM) for 24h and subsequently induced with Dox (1μg/mL) for 48h was analyzed by immunoblotting assays. GAPDH was used as the loading control. (C). Cell viability of iSLK.BAC16 cells treated with increasing doses of clofazimine (CLF) for 24h was analyzed by CellTiter-Glo assays and normalized to mock-treated cells. Results were calculated from n=3 independent experiments and presented as mean ± SEM.

**Figure S5.**
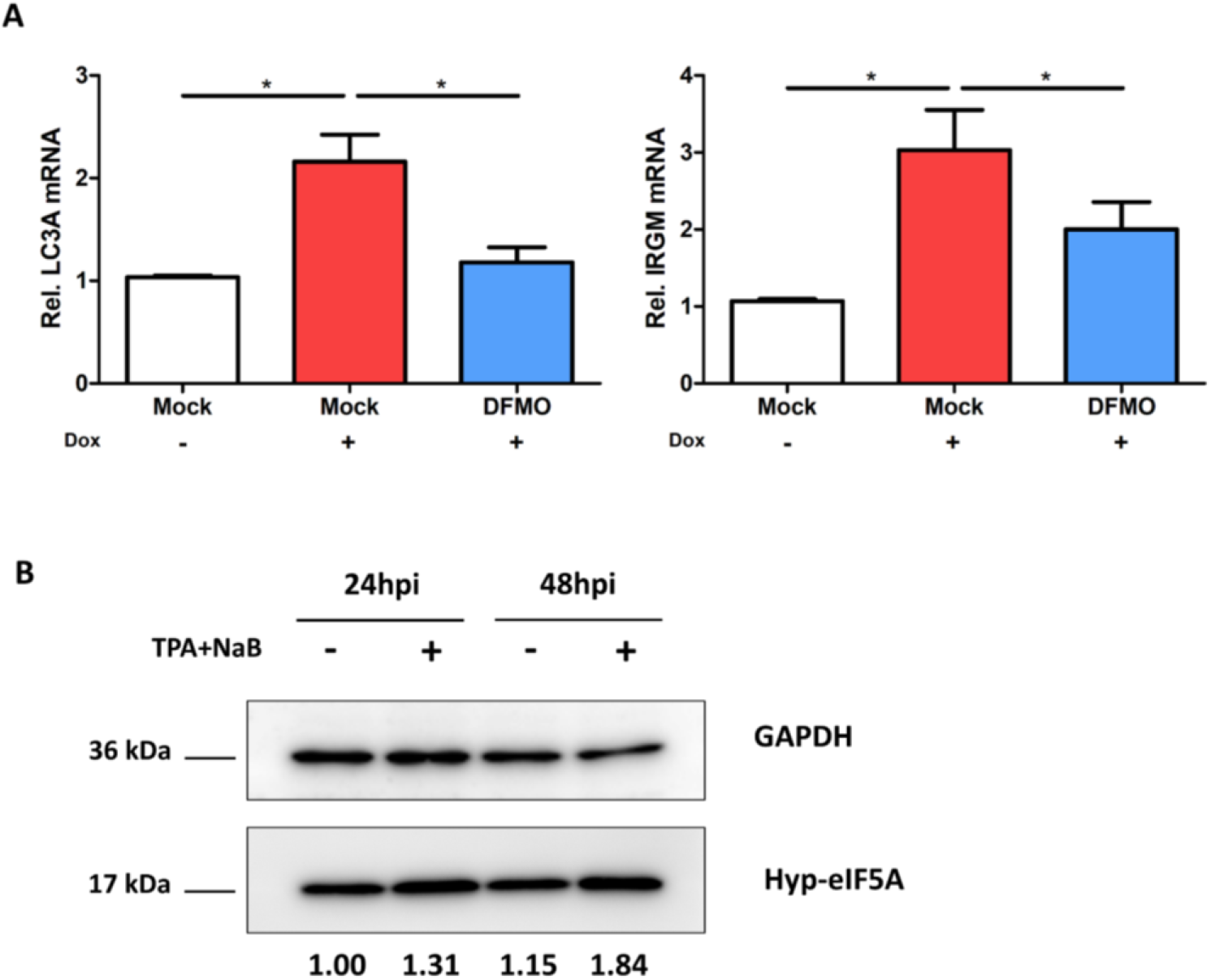
(A). mRNA level of autophagy genes (LC3A, IRGM) in iSLK.BAC16 cells pretreated with DFMO (500μM) for 24h and subsequently induced with Dox (1μg/mL) for 48h was analyzed by RT-qPCR assays and normalized to mock-treated un-induced cells. (B). Protein level of hypusine-eIF5A in HEK293.r219 cells induced with TPA (20ng/mL) + NaB (0.3mM) for 48h was analyzed by immunoblotting assays. GAPDH was used as the loading control. Results were calculated from n=3 independent experiments and presented as mean ± SEM (* p<0.05, two-tailed paired Student t-test).

**Figure S6.**
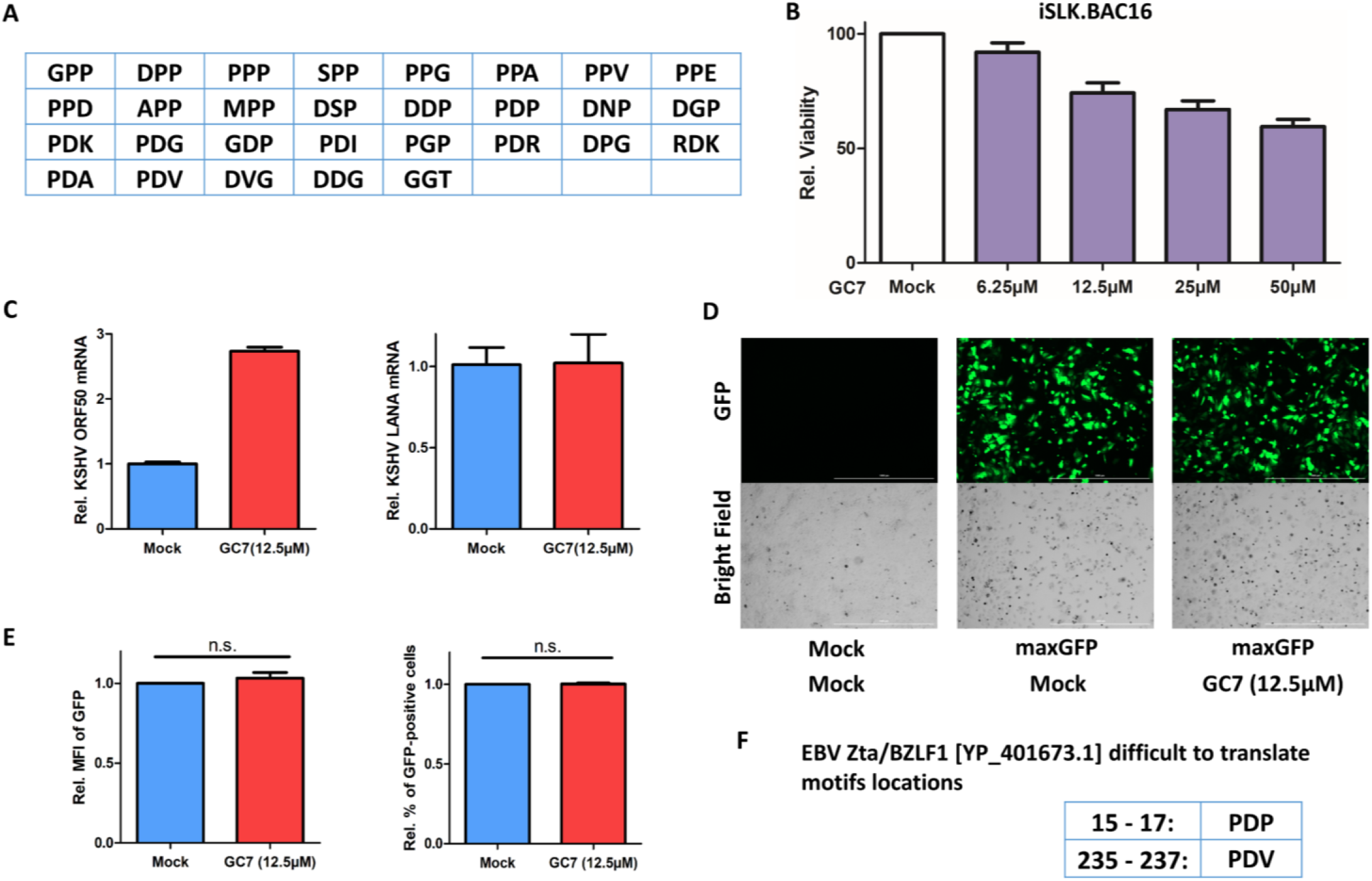
(A). List of hyp-eIF5A-dependent pause motifs reported in Schuller et al. 2017 [17]. (B). Cell viability of iSLK.BAC16 cells treated with increasing doses of GC7 for 24h was analyzed by CellTiter-Glo assays and normalized to mock-treated cells. (C). mRNA level of transfected KSHV ORF50/RTA or ORF73/LANA cDNA in GC7-treated SLK cells was analyzed by RT-qPCR assays and normalized to mock-treated cells. (D, E). SLK cells transfected with pmaxGFP (Lonza) and subsequently treated with GC7 (12.5μM) were visualized by fluorescence imaging (D). These cells were also analyzed by flow cytometry, and relative MFI of GFP expression as well as percentage of GFP-positive cells was determined (E). (F). List of hyp-eIF5A-dependent pause motifs in the coding sequence of EBV Zta protein (YP_401673.1). Results were calculated from n=3 independent experiments and presented as mean ± SEM (ns: not significant, two-tailed paired Student t-test).

**Figure S7.**
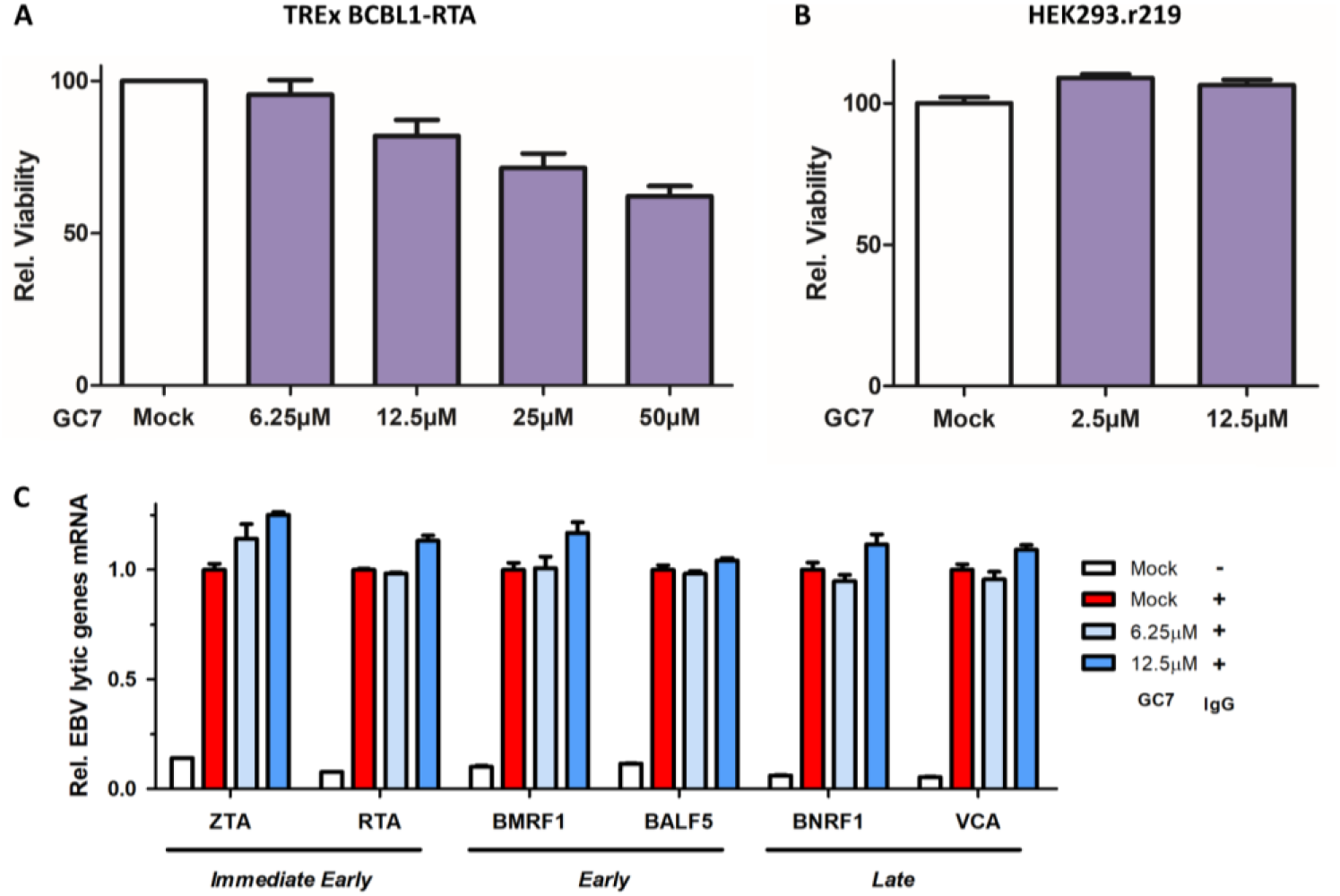
(A, B). Cell viability of TREx BCBL1-RTA (A) and HEK293.r219 (B) cells treated with increasing doses of GC7 for 24h was analyzed by CellTiter-Glo assays and normalized to mock-treated cells. (C) mRNA level of indicated EBV lytic genes in Akata/BX cells pre-treated with GC7 for 24h and subsequently induced with human IgG for 48h was analyzed by RT-qPCR assays and normalized to mock-treated IgG-induced cells. Results were calculated from n=2-3 independent experiments and presented as mean ± SEM.

**Figure S8.**
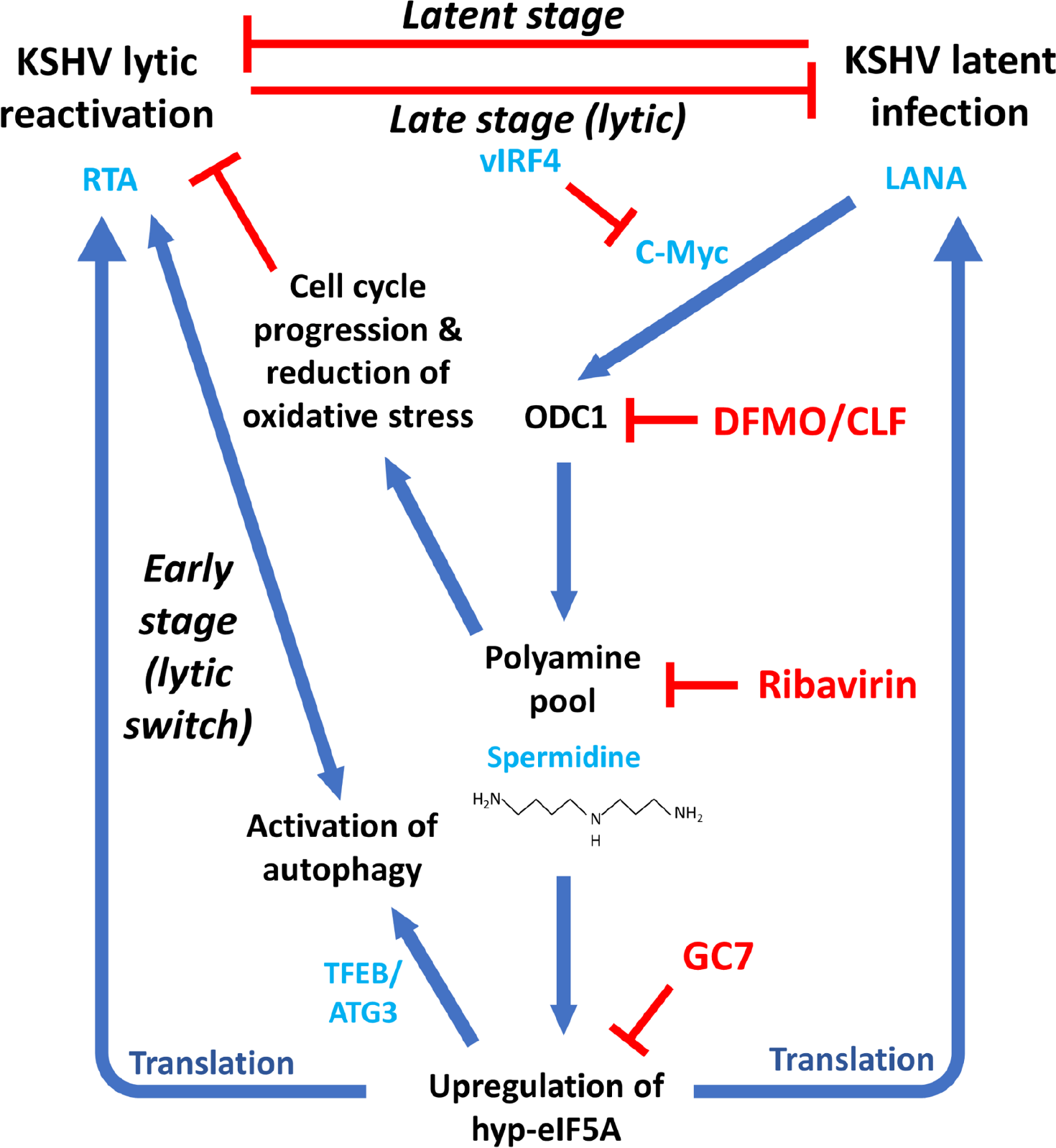
A proposed model illustrates the dynamic and profound interaction of host polyamine biosynthesis and eIF5A hypusination with KSHV infection. At the stage of latent infection, KSHV ORF73/LANA protein induces upregulation of c-Myc that further transcriptionally activates ODC1, which results in the overall increase of intracellular polyamines. However, spermidine is consumed to produce hypusine-eIF5A ensuring the efficient translation of LANA protein required for maintenance of KSHV latency. This results in a positive feedback sustaining activation of the polyamine-hypusine axis. At the early stage of lytic switch, KSHV ORF50/RTA gene starts to be actively transcribed upon certain stimuli, and the constant high level of hypusine-eIF5A not only ensures the efficient translation of RTA protein but also facilitates the activation of autophagy, which overall promotes KSHV lytic reactivation. On the contrary, at the late stage of lytic replication 1061 KSHV RTA protein is expressed up to a sufficient level to suppress LANA protein, leading to decrease of ODC1 and reduction of intracellular polyamines. In addition, other KSHV lytic proteins, such as vIRF4, also contribute to the downregulation of c-Myc, causing further decrease of ODC1 and polyamines. This would generate an adverse effect on cell cycle progression and an increase of oxidative stress, which would be beneficial to KSHV lytic reactivation. Thus, cells are now adjusted to a favorable environment to further promote KSHV lytic replication and viral propagation. KSHV de novo infection likely suppresses ODC1 as well to allow early steps of viral replication. Given to the critical roles of polyamine biosynthesis and eIF5A hypusination in KSHV infection, inhibitors targeting these metabolic pathways would serve as novel antiviral reagents to efficiently block both KSHV latent and lytic replications.

**Table S1.**
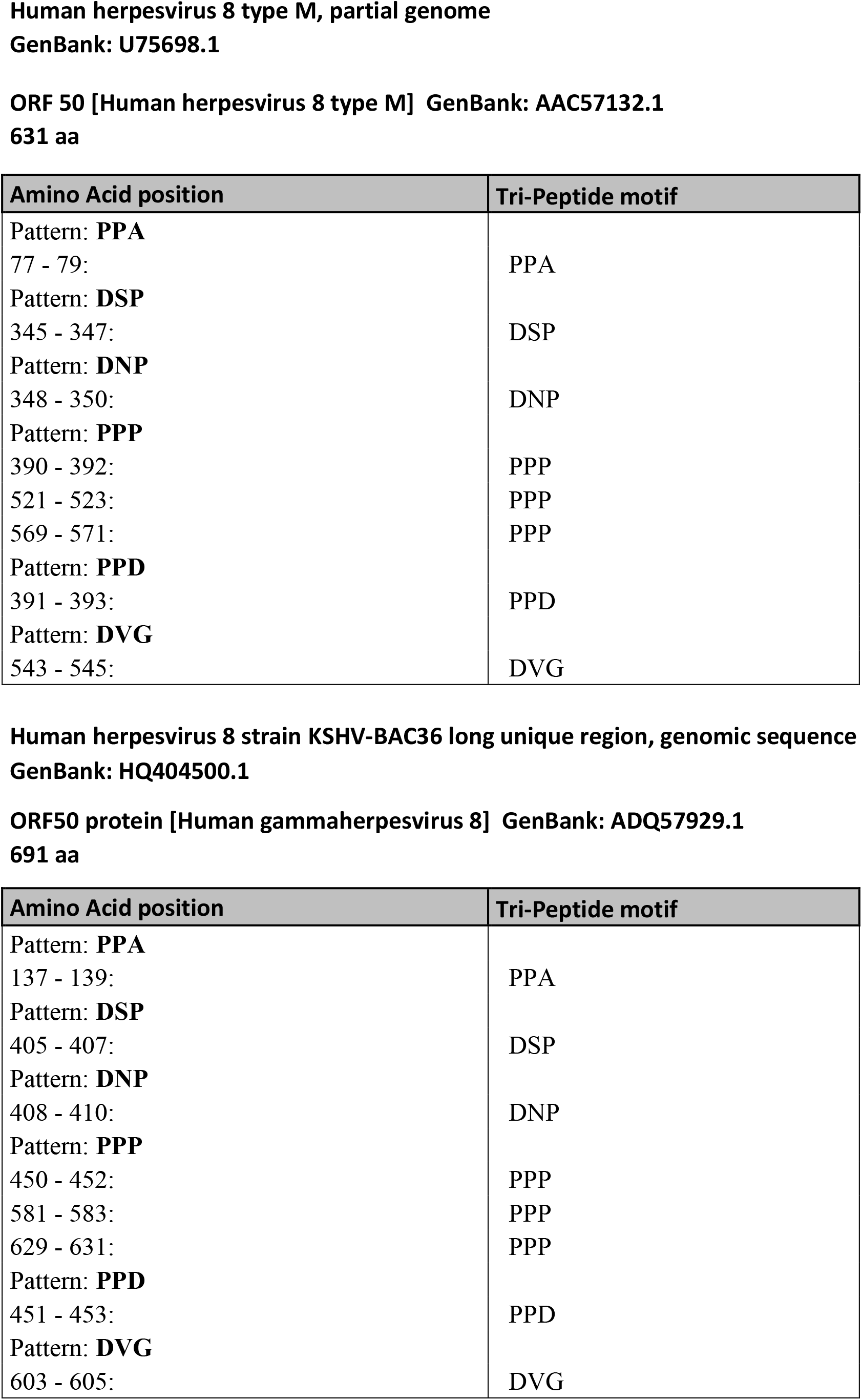
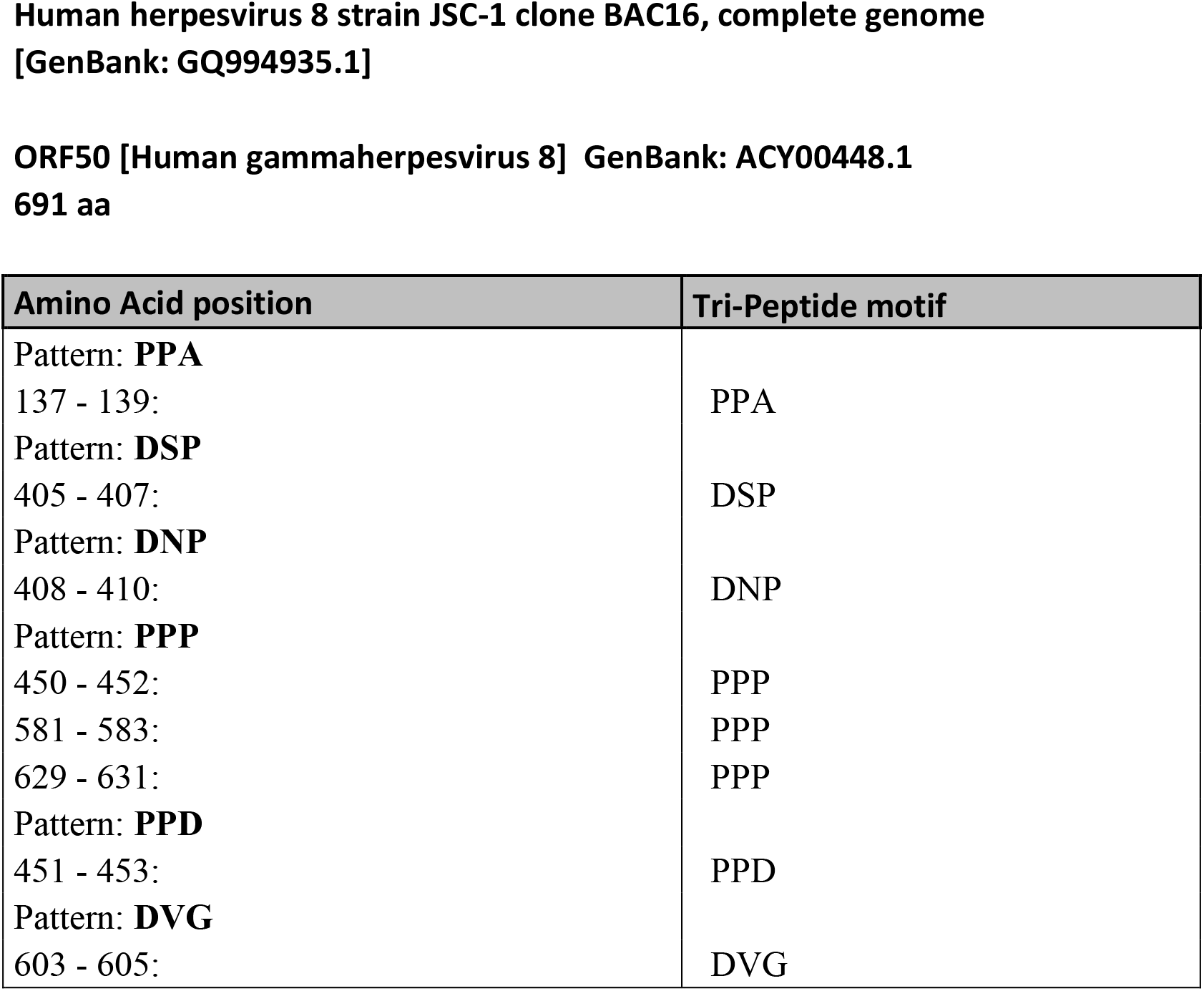
List of hyp-eIF5A-dependent pause motifs in the coding sequence of ORF50/RTA protein from multiple KSHV strains.

**Table S2.**
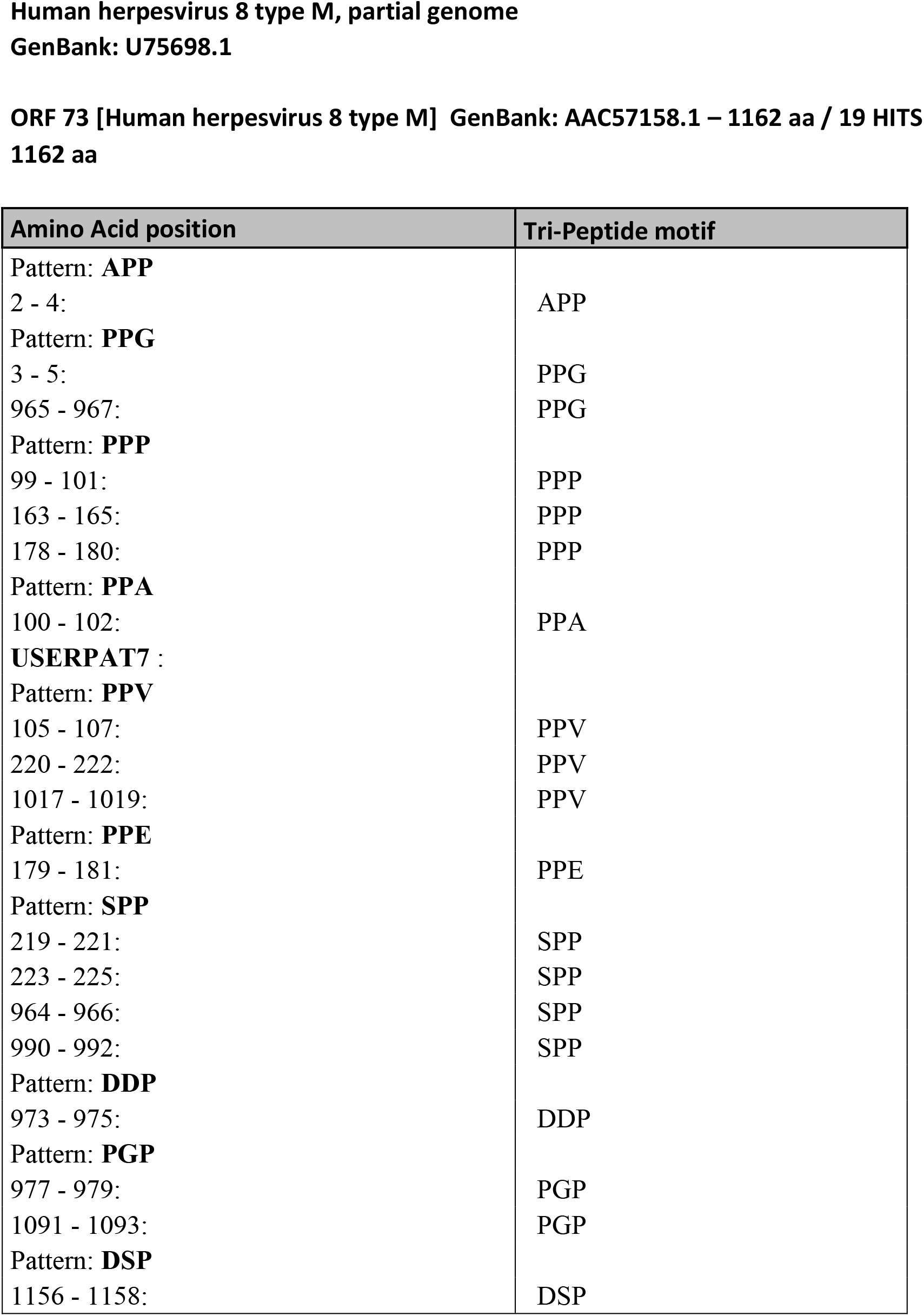
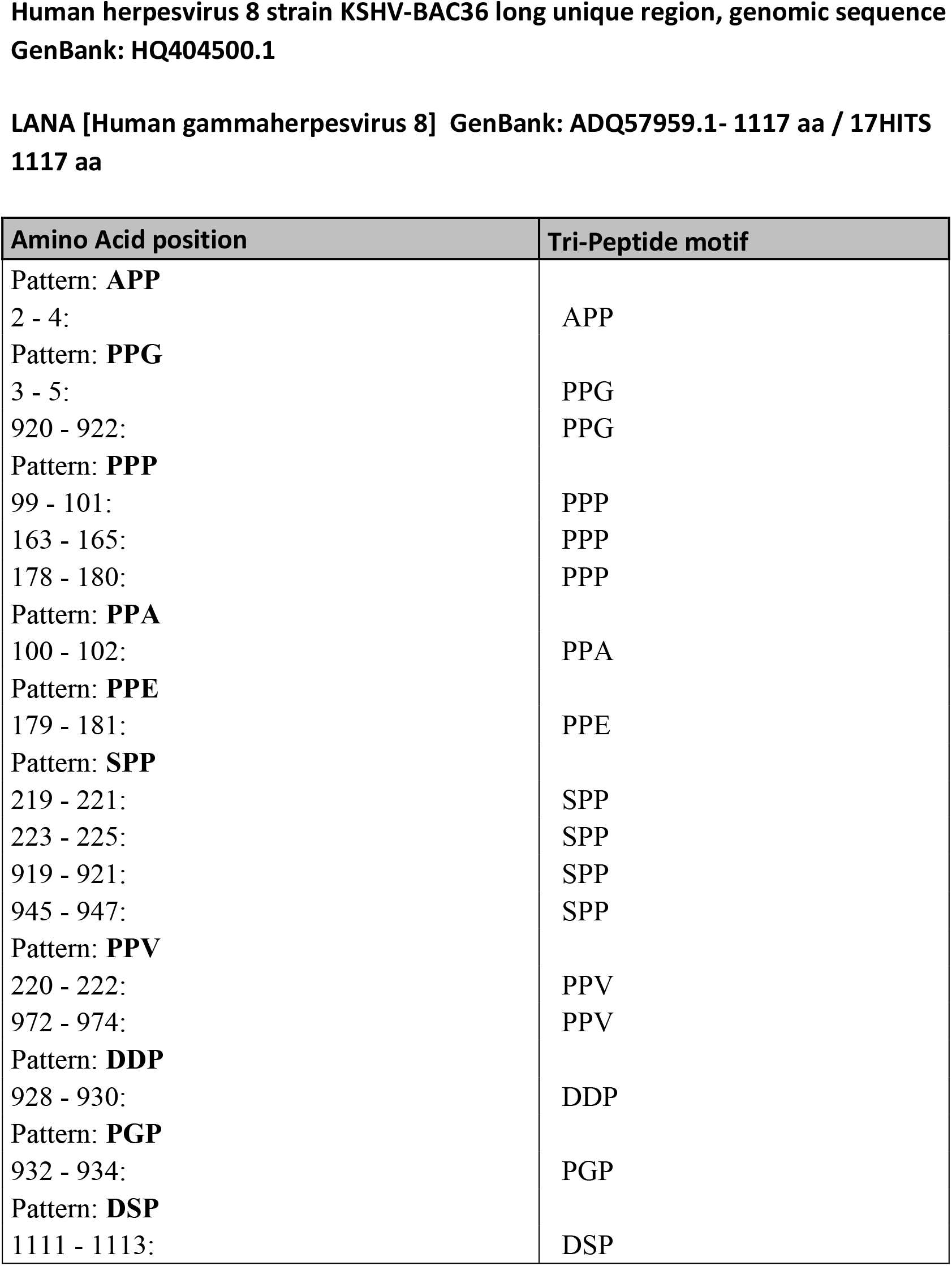
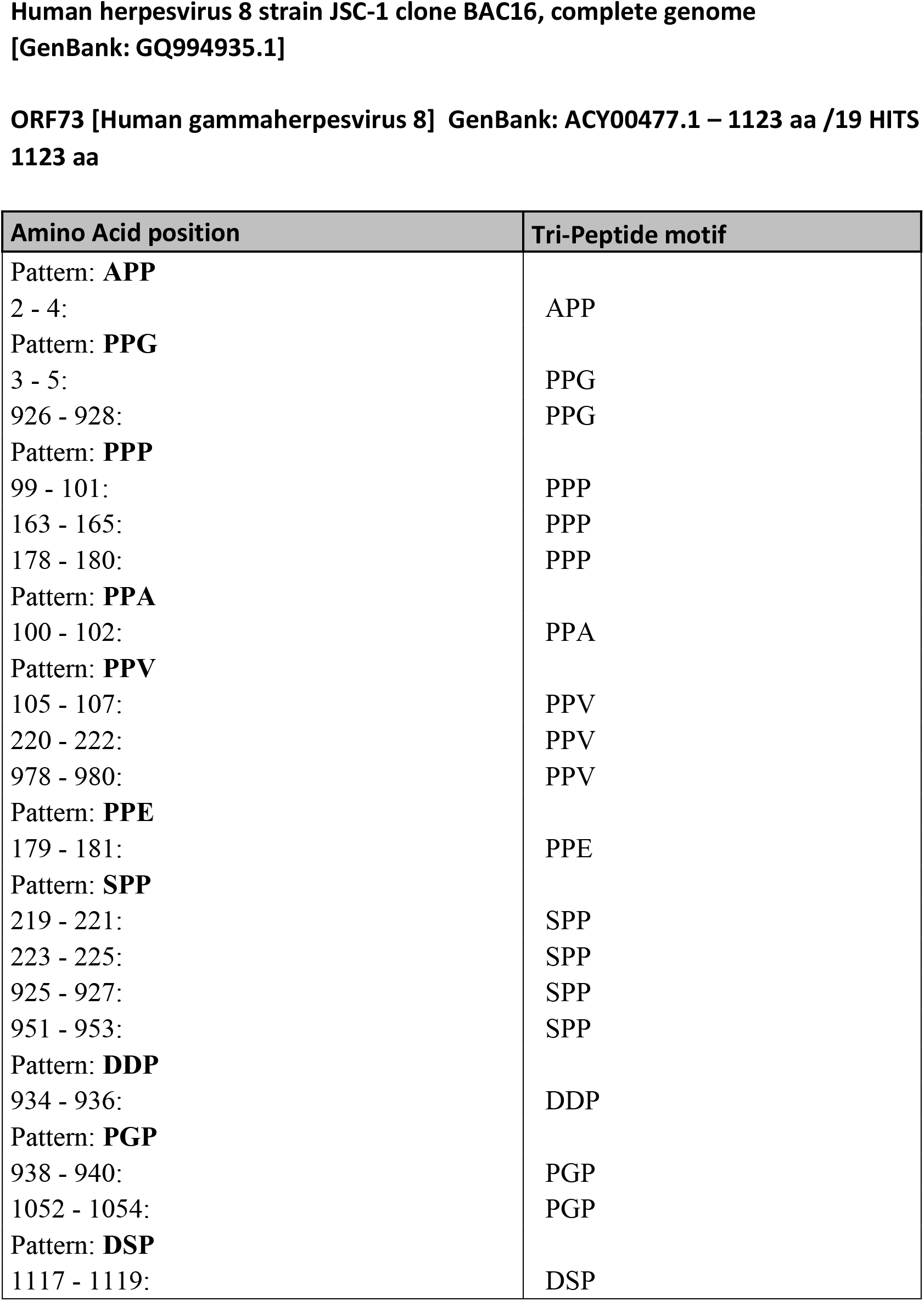
List of hyp-eIF5A-dependent pause motifs in the coding sequence of ORF73/LANA protein from multiple KSHV strains.

**Table S3.**
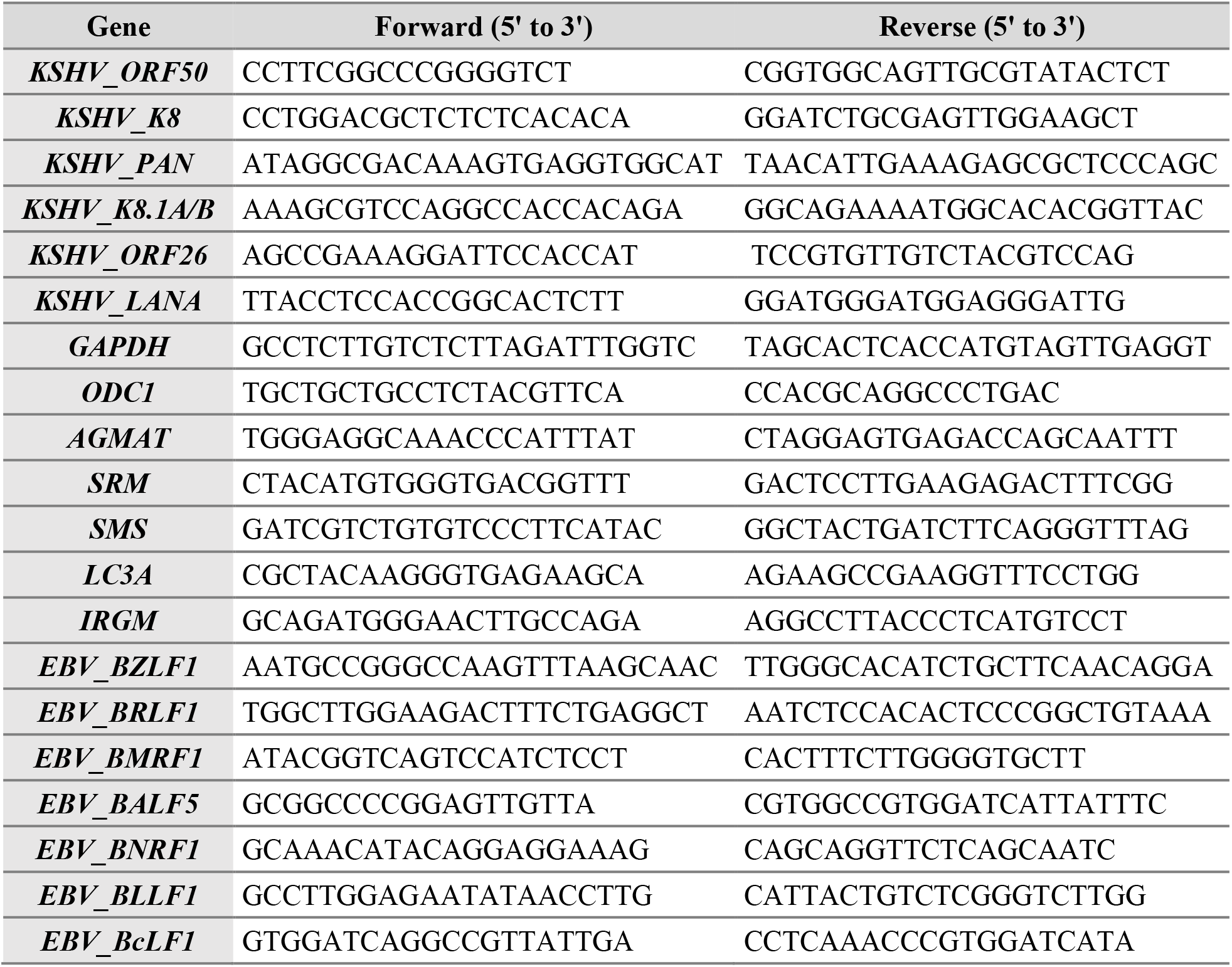
List of PCR primers used in this study

